# Microscopic Chromosomal Structural and Dynamical Origin of Cell Differentiation and Reprogramming

**DOI:** 10.1101/2020.07.16.207662

**Authors:** Xiakun Chu, Jin Wang

## Abstract

As an essential and fundamental process of life, cell development involves large-scale reorganization of the three-dimensional genome architecture, which forms the basis of gene regulation. Here, we develop a landscape-switching model to explore the microscopic chromosomal structural origin of the embryonic stem cell (ESC) differentiation and the somatic cell reprogramming. We show that chromosome structure exhibits significant compartment-switching in the unit of topologically associating domain. We find that the chromosome during differentiation undergoes monotonic compaction with spatial re-positioning of active and inactive chromosomal loci towards the chromosome surface and interior, respectively. In contrast, an over-expanded chromosome, which exhibits universal localization of loci at the chromosomal surface with erasing the structural characteristics formed in the somatic cells, is observed during reprogramming. We suggest an early distinct differentiation pathway from the ESC to the terminally differentiated cell, giving rise to early bifurcation on the Waddington landscape for the ESC differentiation. Our theoretical model including the non-equilibrium effects, draws a picture of the highly irreversible cell differentiation and reprogramming processes, in line with the experiments. The predictions from our model provide a physical understanding of cell differentiation and reprogramming from the chromosomal structural and dynamical perspective and can be tested by future experiments.

## 1 Introduction

Cell is the basic unit of life. Cells have two essential functions. One is the division for reproduction, and the other is differentiation for multi-purpose functionality. Cell differentiation has been suggested to be determined by the underlying gene regulatory network [1]. A fundamental question is how the unique gene expression pattern is progressively established in the specialized cells during differentiation. The roles of epigenetic modifications [2], such as DNA methylation and histone modifications, have been highlighted in modulating the gene expression across different cell types and lineages [3]. As the scaffold for gene expression, genome structure is constantly altered by epigenetic modifications [4]. Therefore, elucidating the microscopic molecular structural origin of cell differentiation at the genomic level is critical for understanding the differentiation mechanism yet still challenging.

Increasing evidence suggests that genome folds into a 3D architecture [5, 6, 7]. Derived from the chromosome conformation capture techniques [8], Hi-C measures the frequency of genomic loci spatially close to each other, resulting in a high-resolution chromosome contact map to infer chromosome structure [9]. The prominent findings of Hi-C show that the chromosome is hierarchically organized through multiple scales from the local self-interacting domains, termed as topologically associating domains (TADs) [10, 11, 12], to the long-range segregation patterns, termed as compartments [9]. TADs are characterized as the functional units of the genome in controlling the gene expressions by restricting the promoters and enhancers within one TAD [13,7,14]. On the other hand, compartments form two types of mutually excluded segregations, referred to as compartment A and B, corresponding to the gene-rich, open active euchromatin and the gene-poor, closed repressive heterochromatin, respectively [9,15,16]. TADs and compartments are critical genome structural features that are found to play important roles in many biological processes, including DNA replication [17], transcription [7] and gene regulation [18, 19, 20].

In the context of Waddington’s epigenetic landscape [21], the cell differentiation process can be metaphorically referred to as an iconic picture, where a ball rolls down on a surface (landscape) from the top to the lowest point. The guidance of the ball in seeking stable positions, which are the differentiated cell states, is controlled by the landscape that is shaped by the epigenetic mechanism. Genome structure, which is in intimate relation to epigenetic modifications, has been found to undergo high-order structural rearrangement during differentiation [22], in order to accommodate the required gene expression pattern possibly through modulating the accessibility of the genomic loci to transcription factors. Therefore, cell differentiation should be associated with a landscape at the microscopic molecular level that describes the genome structural dynamics, as a reminiscence of the one widely used in protein folding [23]. The information of this landscape can provide a physical understanding of genome dynamics and microscopic molecular structural origin for cell differentiation, giving insights into the underlying mechanism. Therefore, such structural landscape at the genomic level can serve as the physical and structural basis for understanding and quantifying the Waddington landscape at the gene expression level for cell differentiation and development. However, exploring such structural landscape in practice is very challenging due to the lack of precise determination of the chromosome structure and dynamics, as well as the non-equilibrium nature of cell differentiation distinctly different from protein folding. A natural initial step towards this ultimate goal is to quantify the chromosomal structural rearrangement during cell differentiation.

Embryonic stem cells (ESCs) are pluripotent cells and can differentiate to the somatic cells during cell development. On the other hand, a somatic cell can be reprogrammed back to the induced pluripotent stem cell (iPSC), which genetically behaves like the ESC with high pluripotency [24, 25]. Cell reprogramming, induced by ectopic expression of the key transcriptional factors, erases the epigenetic modifications established in the differentiated cell [26], and results in the reorganization of chromosome structure into an ESC-like form [27]. Understanding how the chromosome undergoes the structural rearrangement during reprogramming has important implications in tissue engineering for developing new approaches to improve the reprogramming efficiency, however, is currently not well addressed. The understanding of cell reprogramming at chromosome structural level can complete our knowledge of the forward and backward cell development processes, however, is still lacking.

Herein, we developed a non-equilibrium landscape-switching model to investigate chromosome dynamics during cell differentiation and reprogramming. The key element that controls and induces the differentiation and reprogramming was modeled as an energy excitation that triggers the processes for switching of chromosome structural dynamics between the two landscapes, representing the ESC and the terminally differentiated cell, followed by the subsequent conformational relaxation. The model allowed us to accumulate a statistically sufficient number of differentiation and reprogramming trajectories and perform the subsequent chromosome structural analysis. We found that the chromosome shows compartment-switching in the units of relatively stable TADs during both the differentiation and reprogramming processes. Chromosome in differentiation undergoes gradually monotonic compaction with spatial re-positioning of active and inactive genomic loci, while an over-expanded chromosome structure with universal localization of the loci at the chromosome surface appears in reprogramming. Further analysis of the differentiation process exhibited a distinct pathway that bifurcates at the quite early developmental stages to the destined differentiated cell. Our model made extensive predictions that have suggested a chromosomal-level structural and dynamical picture of cell differentiation and reprogramming.

## 2 Results

### 2.1 Chromosome dynamics during cell differentiation and reprogramming

We developed a non-equilibrium excitation-relaxation energy landscape-switching model to investigate the chromosomal structural rearrangement during the differentiation of the ESC to the somatic cell and the reprogramming of the somatic cell to the iPSC. The model is similar to the one used for studying the cell-cycle chromosome dynamics in our previous study [28]. To simplify the processes, here we used the ESC to replace the iPSC as the destination for reprogramming. This is a reasonable approximation based on the recent evidence that the human ESC and iPSC are equivalent in terms of gene expression patterns [29]. On the other hand, we used the IMR90 as the somatic cell in our study. IMR90 is the terminally differentiated human fetal lung fibroblast cell and has been proved capable of reprogramming to the iPSC by various transcriptional factors [30].

We focused on the long arm of chromosome 14 (20.5Mb-106.1Mb). With a resolution of 100 kb, the chromosome segment was reduced into a coarse-grained polymer representation with 857 beads. The Hi-C maps and compartment profiles of the chromosome segment at the ESC and IMR90 show distinct differences (Figure 1), indicating that significant structural rearrangement should occur during the transitions between these two cell states. We first used the maximum entropy principle to implement Hi-C data to a generic polymer model as a minimally biased experimental restraint [31, 32]. This approach provides two effective energy landscapes that generate the experimentally consistent chromosome structural ensembles and kinetically govern the chromosome dynamics at the ESC and IMR90, respectively (Figure S1-S2 in SI) [33, 34]. It is worth noting that chromosome dynamics at both the ESC and IMR90 are out-of-equilibrium [35, 36, 33]. However, an effective energy landscape was demonstrated to be capable of describing the behaviors of non-equilibrium systems in certain cases not too far from equilibrium [37, 38, 39]. Then we triggered the differentiation/reprogramming process by switching between these two effective landscapes. Finally, the chromosome was relaxed under the new effective landscape. In practice, we termed the switching and relaxation from the ESC to the IMR90 landscape as a differentiation process and from the IMR90 to the ESC landscape as a reprogramming process, respectively. Such switching can be regarded as an energy excitation that activates the system to trigger the escape from the local minima on the energy landscape and achieves the transition to a new one. This excitation requires energy input or cost, which is provided by the underlying molecular processes (e. g., ATP hydrolysis). The net energy input or cost leads to the detailed balance breaking of the system, resulting in a non-equilibrium process. Therefore, the model captured the essence of cell differentiation and reprogramming processes and can provide insights for the underlying non-equilibrium dynamics. Further details of the model can be found in ‘‘**Model and Simulation Section**” and our previous study [28].

**Figure 1:**
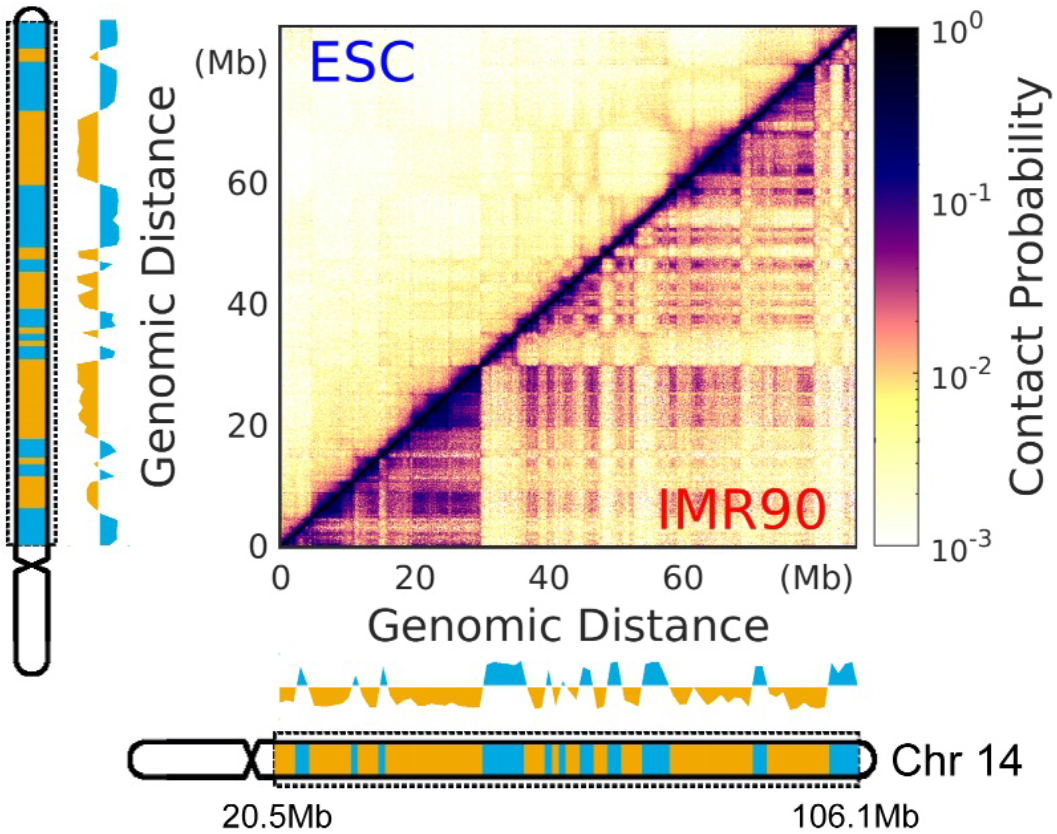
Structural differences of the chromosome segment in the ESC and IMR90. The chromosome segment focused in our study is from the long arm of chromosome 14 with a range of 20.5-106.1 Mb. Hi-C maps of the chromosome for ESC *(Left)* and IMR90 *(Right)* are normalized from 0 to 1. The compartment profiles are shown close to the Hi-C maps and colored by the compartment states with A and B corresponding to blue and yellow, respectively. The ideograms of chromosome 14 are shown. The bands within the chromosome segment used in our study (denoted by the dashed rectangle) are colored by the compartment states. Both the compartment profiles and ideograms are aligned to the corresponding Hi-C maps *(Left* as ESC and *Bottom* as IMR90).

We accumulated a statistically sufficient number of chromosomal structural rearrangement pathways during the ESC differentiation and the IMR90 reprogramming, and then calculated the contact probability map of the chromosome at each time frame, referring to as the evolution of the Hi-C heat map in differentiation and reprogramming (Figure 2). We quantified the similarity of the Hi-C maps by a sum of the differences of the pairwise contact probability 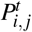 (for chromosomal loci *i* and *j*) between the time frames *t = I* and *t = J,* and then obtained a time-dependent matrix Δ*P^I,J^* (Figure 2A and 2F). These matrices clearly show that chromosomal structural rearrangement during both differentiation and reprogramming undergo stepwise changes. Since chromosome structure is organized at multiple levels (e. g., TADs and compartments), we further calculated Δ*P^I,J^* at local and non-local contact ranges based on the genomic distance between the involved contact pair of chromosomal loci *i* and *j*. Hierarchical clustering of matrices Δ*P^I,J^* exhibits two apparent large clusters that can roughly group the trajectories into two steps or stages in the differentiation and reprogramming processes (Figure 2A and 2F), indicated as blue and red linkages on the hierarchical trees. Moreover, we observed different time points for separating these two linkages at local and non-local contacts, implying that the local and non-local contacts in the chromosome evolve asynchronously. During differentiation (Figure 2A), the chromosome appears to form the destined local contacts faster than the non-local ones, while the local and non-local structural changes in the chromosome are more synchronous during reprogramming (Figure 2F). We then calculated the contact probability *P*(*s*) versus the genomic distance (*s*) in the chromosome and we see the slope *a* in the relation of *P*(*s*) ~ *s^a^* gradually changes from −1.5 to −1.0 in differentiation and −1.0 to −1.5 in reprogramming (Figure 2B and 2H, Figure S7A, Figure S7B, S8A and S8B). This is consistent with the consensus that the chromosome in the human somatic cell behaves as a fractal-like globule [40] (Figure S4) and is more loosely formed in the ESC (Figure S3).

**Figure 2:**
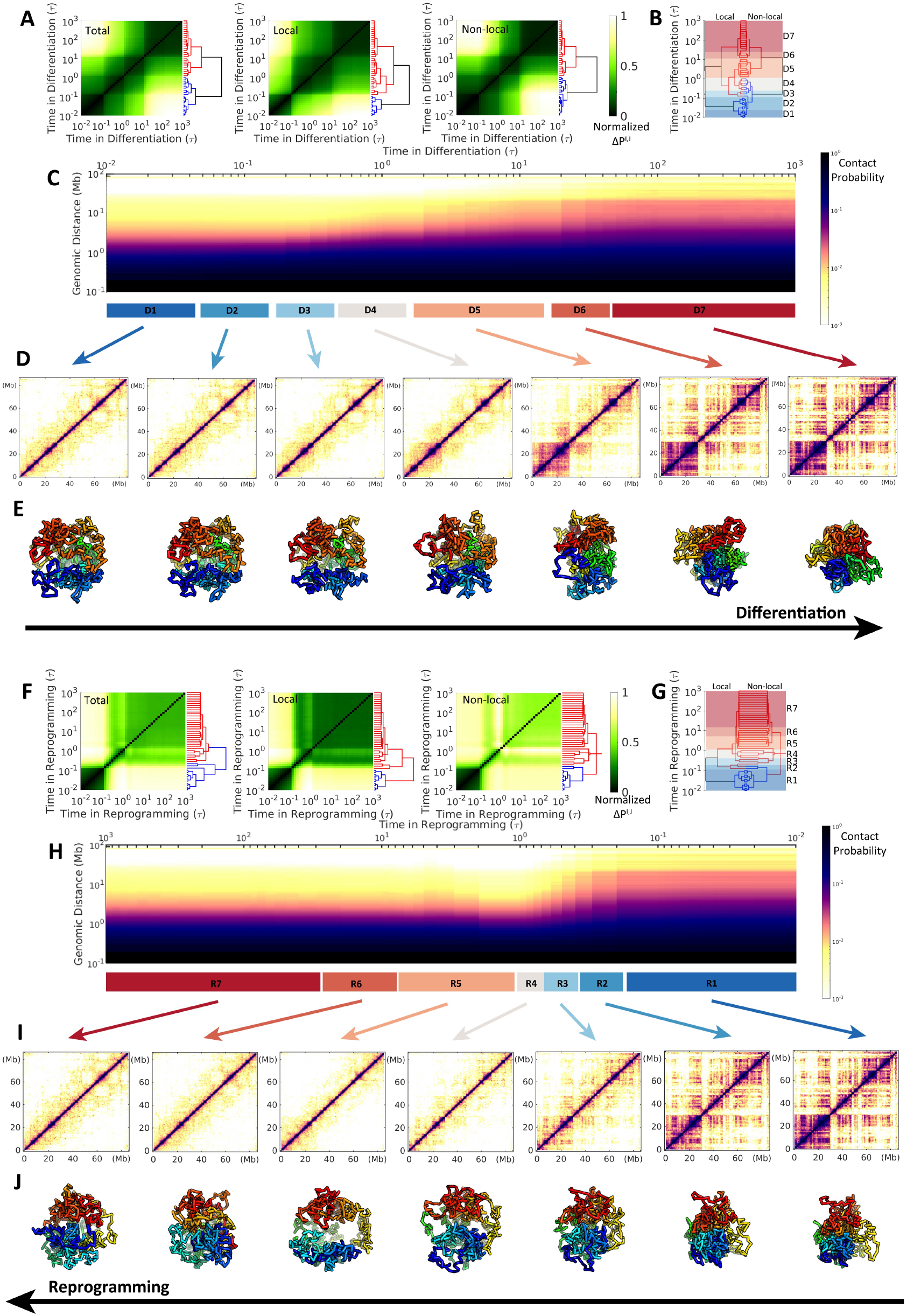
Chromosomal structural rearrangement during the ESC differentiation *(Upper)* and the IMR90 reprogramming (Lower). (A) Differences and hierarchical clustering of contact probability maps of the chromosome among each time frame *t = I, J* during differentiation varied by total, local (< 2Mb) and non-local (> 2Mb) contact ranges. The contact map difference Δ*P^I,J^* is calculated by: 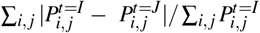, where 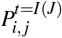 is the contact probability between chromosomal loci *i* and *j* at the differentiation time *t = I* or *J*. Δ*P^I,J^* is further normalized for achieving a better visualization. (B) Reduced 7 stages (“D1”-“D7”) for the differentiation process based on the combination and comparison of the hierarchical clustering trees of local and non-local *ΔP^I,J^* established in (A). The chromosome within one stage thus possesses a relatively similar contact probability map. (C) The change of the contact probability *P*(*s*) versus genomic distance *s* in the chromosome during differentiation with the 7 stages indicated at the bottom. (D) Hi-C heat (contact probability) maps of chromosome for the 7 stages during differentiation. (E) Representative structures of the chromosome for the 7 stages during differentiation. (F-J) are similar with (A-E) except for the process of IMR90 reprogramming to ESC. Another reduced 7 stages during the reprogramming (“R1”-“R7”) are determined in (G).

We further reduced the complexity of differentiation and reprogramming by respectively coarse-graining the processes into 7 stages based on hierarchical clustering of chromosome contact map similarity Δ*P^I,J^* (Figure 2B and 2G). This was done by directly comparing the hierarchical trees for local and non-local Δ*P^I,J^* and then sequentially and gradually grouping the time-evolving states that have high similarity at both local and non-local ranges from the contact map perspective (states within one stage are within the same cluster in terms of similar formed contacts). Chromosomes at these stages are shown with Hi-C heat maps and representative structures (Figure 2D, 2E and 2I, 2J). The plaid pattern on the chromosome contact probability map in the IMR90 is gradually gained (lost) during the differentiation (reprogramming) process along with the shape changing from large to small (small to large) globules. Careful examination on reprogramming can detect possible over-expanded chromosome structures at stages R4 and R5, where P(s) is over-decreased under that in the ESC (Figure 2H and Figure S8A). In contrast, the chromosome undergoes monotonic compaction during differentiation (Figure 2C and Figure S7A). These results clearly show that the chromosome adapts large-scale structural rearrangement during cell differentiation and reprogramming, and these two processes do not share the same pathway at the chromosomal level.

### 2.2 Chromosomal structural rearrangements in the units of TADs

Since the TAD is regarded as the structural and functional unit of the chromosome [41, 17, 7] and the compartment spatially segregates the active and inactive genomic loci from functional perspective [9], here we examined chromosomal structural rearrangement in terms of TADs and compartments during differentiation and reprogramming. Compartment profiles were calculated along all the loci in the chromosome segment, and the reduced 7 stages in differentiation and reprogramming are presented (Figure 3A and 3G). We can see that the chromosome undergoes apparent compartment-switching during both of the two processes. In detail, differentiation exhibits a more substantial degree of switching from compartment A to B (blue to yellow) than that from compartment B to A (yellow to blue) (Figure 3A). This is a sign of more regions switching from active to inactive loci, consistent with the fact that the ESC possesses the most expressive euchromatin in order to execute the highest pluripotency [42, 22]. This is oppositely found during cell reprogramming (Figure 3G). In addition, the most significant compartment-switching during differentiation occurs at stage D5 (Figure 3A), which appears to be after 2τ, much slower than the structural rearrangements in TADs (Figure 3F). Since compartment forms at a longer range (beyond 5Mb) than TAD does, this shows again that local chromosome structural arrangement occurs faster than nonlocal one does. On the other hand, we observed quite different compartment-switching behaviors in the chromosome during reprogramming, which is not a simple reversal of that during differentiation. The compartment profiles appear to proceed with gradual change during reprogramming from stages R1 to R4 (Figure 3G). At stage R5, a large proportion of switching in compartment profiles occurs with extensive regions showing that the associated compartment profiles are different from those in either the ESC or IMR90 (e. g., at 6-22Mb). This implies an aberrant structural rearrangement in the chromosome. We note that the chromosome in R5 exhibits an over-expanded configuration. Such configuration has completely disturbed the chromosome compartment formation reserved in the ESC and IMR90. This is also evidenced by the enhanced contact probability map (Figure S9), which essentially determines the compartment profiles [43].

**Figure 3:**
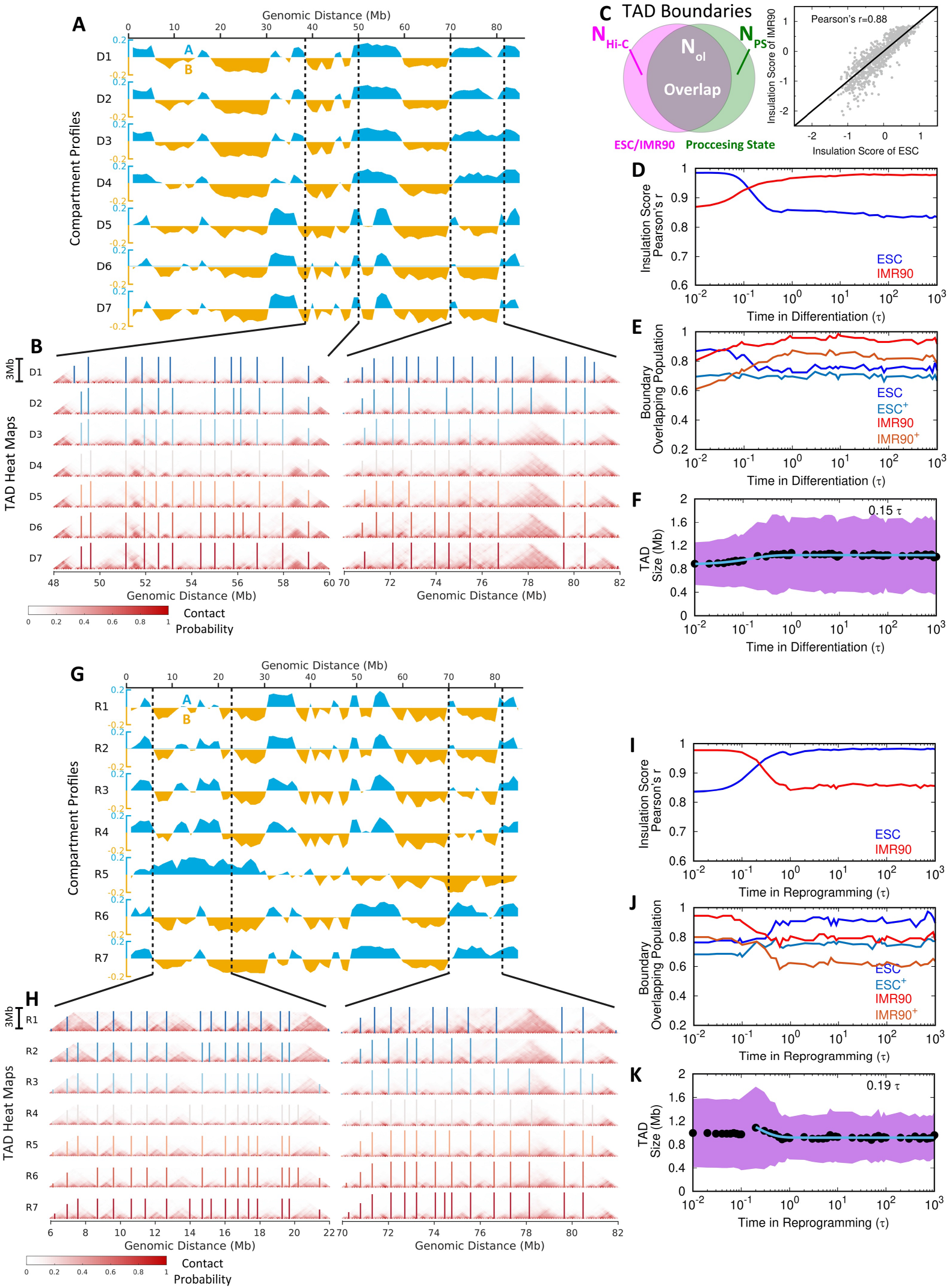
Extensive switching of compartments A/B and conserved TADs in the chromosome during differentiation and reprogramming. (A) Compartment profiles of the chromosome for the 7 stages during differentiation. The values of compartment profiles are defined as the first principal component (PC1) obtained by the principal component analysis (PCA) of the contact probability maps (Details in “**Model and Simulation Section**”). Compartments A and B are shown blue (positive) and yellow (negative) along with the whole range of the chromosome segment, respectively. (B) Hi-C heat (contact probability) maps of the two local regions of the chromosome that have compartment-switching indicated in (A) for illustrating the structural changes of TADs at the 7 stages during differentiation. The boundaries of TADs are determined by the insulation score [44] and shown with vertical lines in the heat maps. (C) An illustration of TAD boundary overlapping between Hi-C data (ESC or IMR90) and simulation processing state during differentiation and reprogramming (*Left*). The numbers of TAD boundaries detected by insulation score for experimental Hi-C (ESC or IMR90), simulation processing state, and the overlap between processing state and Hi-C (ESC or IMR90) are denoted as 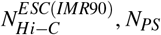, *N_PS_* and 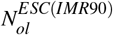, respectively. Insulation scores obtained from the experimental Hi-C data show high correlations between the ESC and IMR90 *(Right).* (D) The change of Pearson’s correlation coefficient of the insulation score obtained from the simulation to the one from experimental Hi-C of the ESC and IMR90 during differentiation. (E) The change of TAD boundary overlapping population from the simulation processing state to the Hi-C of the ESC and IMR90 (calculated by 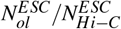 and 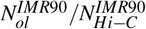) and from the Hi-C of the ESC and IMR90 to the simulation processing state (indicated by superscript ^+^ and calculated by 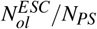 and 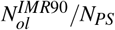) during differentiation. (F) The change of the TAD size during differentiation. The data are fitted to exponential function with the relaxation time shown. Shadow region represents the standard deviations of the TAD size at the corresponding time. (G-K) are similar to (A, B, D, E, F) except for reprogramming. The time shown in (K) is the relaxation time after passing the over-expansion stage.

On the other hand, we observed very similar Hi-C heat maps at local regions in the chromosome during differentiation and reprogramming, implying that the structures of TADs are unaltered during these two processes (Figure 3B and 3H). To quantitatively investigate the structural change of TAD, we firstly used the insulation score, proposed by Crane et al. [44], to define the TAD boundary (Figure S1C and S2C). We see that most of the TAD boundaries are conserved during differentiation and reprogramming (bars in Figure 3B and 3H). This was also observed in stages D5 and R5, where the compartment profiles of the chromosome have significantly changed. Since the boundaries of TADs in the ESC and IMR90 are strongly overlapped (Figure 3C), the differentiation and reprogramming processes do not change TADs significantly in a way that would deviate from the destined states. This is also quantitatively confirmed by the observation that the correlation between the insulation score and the TAD boundary overlapping population to the destined state (IMR90 in differentiation and ESC in reprogramming) increases monotonically as the process proceeds (Figure 3D, 3E, 3I and 3J).

Therefore, our findings have reached a conclusion that chromosomal structural rearrangements during differentiation and reprogramming undergo changes in the unit of TADs. This is consistent with the fact that TADs are conserved structural units across different cell types [10]. However, through the detailed analysis, we observed that the size of TAD slightly increases and decreases in differentiation and reprogramming, respectively (Figure 3F and 3K). Interestingly, the over-expanded chromosome during reprogramming possesses a larger average TAD size than that in both the ESC and IMR90, followed by a quick relaxation process to the destined state. This implies that expanding of TADs also occurs within the over-expanded chromosome. Overall, we conclude that the chromosome rearranges its structure mostly through the compartment-switching without significantly disturbing the local TAD formation.

### 2.3 Identification of cell differentiation and reprogramming pathways and irreversibility of chromosomal structural rearrangements

To investigate the chromosomal structural rearrangements during differentiation and reprogramming, we quantified the transition pathways and projected them onto different structural order parameters. It is shown that the pathways are highly stochastic during both differentiation and reprogramming (Figure 4). We first examined from the geometrical perspective by using the structural extension at the three principal axes (PA) (Figure 4A, *Left* panel). We show that the chromosome compacts its configuration via a two-step scenario with initially shortening along its shortest PA (PA3) followed by compressing along with the longest PA (PA1) during differentiation. Such anisotropic compaction can also be observed by projecting the pathways onto the radius of gyration (*R_g_*) and an anisotropy measure, the aspheric quantity △ calculated using the inertia tensor [45]. Deviation of Δ from 0, which corresponds to a perfect sphere, measures the extent of anisotropy. As shown in Figure 4B, *Left* panel, *R_g_* decreases with △ increasing first and then followed by decreasing due to the anisotropic compaction. Interestingly, the chromosome in reprogramming apparently undergoes distinct pathways, which are not the reversal of those in differentiation. The pathways projected on PA1 and PA3 indicate isotropic expansions during the IMR90 reprogramming to the ESC (Figure 4A, *Middle* panel). In particular, there are chromosome structures that exhibit bigger sizes than those in the ESC being observed, as evidence of over-expansion. Such over-expansion followed by slight compaction in the chromosome during reprogramming is also confirmed from the pathways projected onto *R_g_* and Δ (Figure 4B, *Middle* panel).

**Figure 4:**
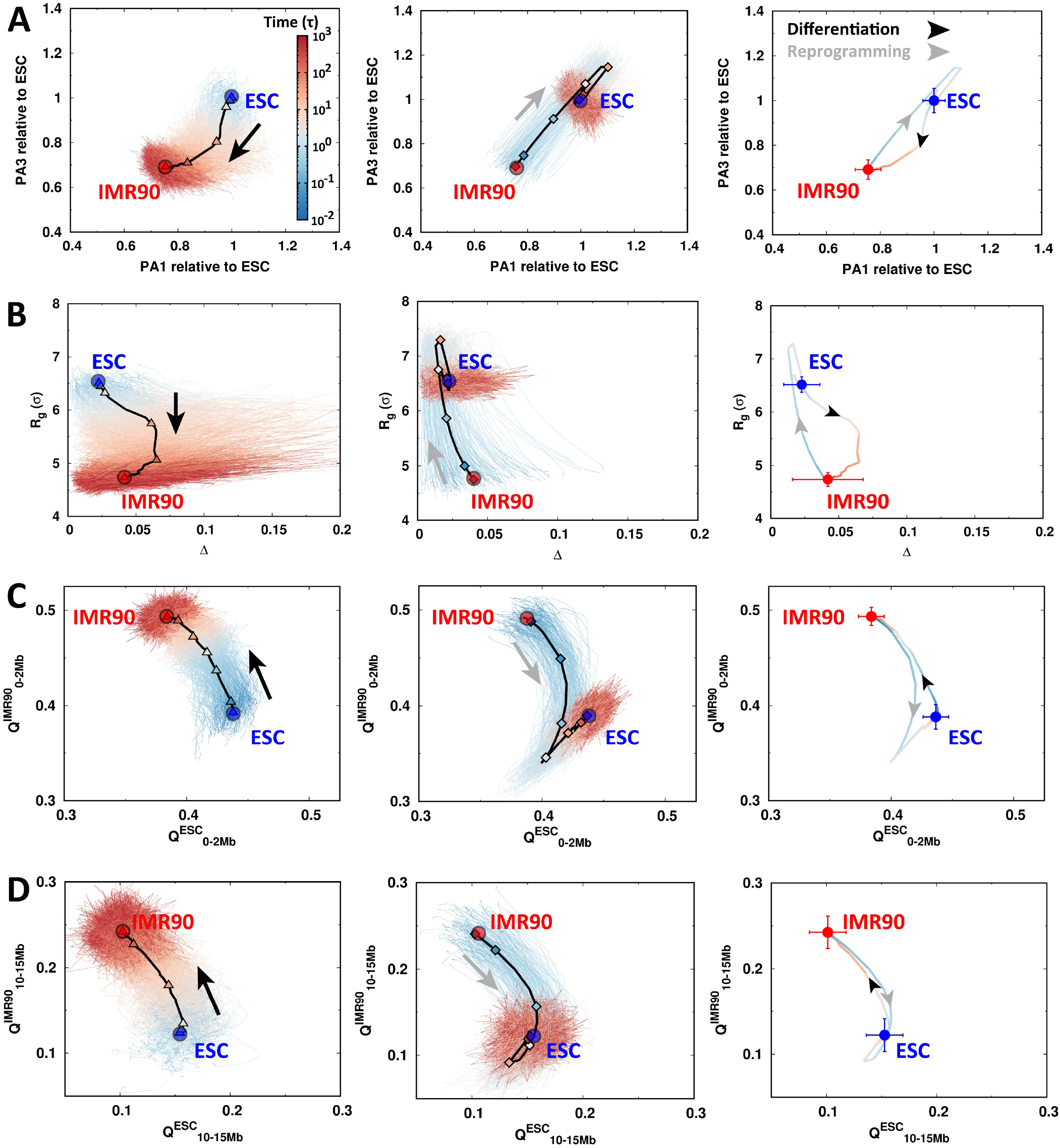
The pathways of chromosomal structural rearrangement during differentiation and reprogramming. The pathways are projected onto the order parameters of the chromosome with all trajectories presented for differentiation (Left) and reprogramming (Middle). The trajectories are colored by time in the logarithmic scale. The 7 stages are indicated by triangles (“D1”-“D7” in differentiation, *Left)* and diamonds (“R1”-“R7” in reprogramming, *Right)* on the averaged pathways shown as black lines. The averaged pathways for differentiation and reprogramming are additionally shown and colored by time (*Right*). The ESC and IMR90 are placed as the blue and red points, respectively. The arrows indicate the directions of differentiation (black) and reprogramming (grey). The pathways are projected onto (A) the extensions of the longest and the shortest principal axes (PA1 and PA3), (B) the radius of gyration (*R_g_*) and the aspheric quantity (Δ), contact similarity in terms of the fraction of native contact *Q* to ESC and IMR90 at (C) local range (0-2Mb), and (D) long-range (10-15Mb).

We then used the fraction of “native” contacts *Q*, which was widely used in protein folding [46], as the order parameter. We practically used the averaged pairwise distances obtained from the maximum entropy principle simulations for the ESC and IMR90 as the references for native distances to calculate *Q* (Figure S3 and S4). Furthermore, we divided *Q* into different categories varied by genomic distances between the contacting pairs. From Figure 4C and 4D (*Left* panels), we can see that during differentiation, the chromosome gradually loses and gains the contacts formed in the ESC and IMR90, respectively. The local contacts (0-2Mb) form faster than non-local ones (10-15Mb) at the early stages of the differentiation process. On the other hand, we observed that the over-expansion occurs in both local and non-local contacts of the chromosome during reprogramming, shown by a simultaneous decrease of *Q^ESC^* and *Q^IMR90^* during reprogramming (Figure 4C and 4D, *Right* panels). Intriguingly, the most significant over-expansion in the chromosome was observed at different stages at local (R4) and non-local contacts (R5). This implies that the chromosome has different expanding rates at different structural levels. By examining the average pathways for differentiation and reprogramming, we found that the two pathways never overlap, indicating that the chromosomal structural rearrangements in these two processes are highly irreversible (Figure 4, *Right* panels).

### 2.4 Rearrangements and transformations of chromosome compartmental structures during cell differentiation and reprogrmming

The transcriptional activity has been found to be strongly correlated with the spatial distribution of the gene regions [47,48]. To see how the chromosome dynamically manages the spatial re-positioning of active and inactive loci, we calculated the radial density of chromosomal loci and examined its change during differentiation and reprogramming (Figure 5). Consistent with previous simulations [49, 33, 50] and experiments [16], we found that in the IMR90, loci in compartment A tends to lie closer to the surface of the chromosome than the ones in compartment B (Figure 5A, red lines in *Middle* and *Right* panels). In contrast, there is not much difference on the spatial positioning of loci in compartment A and B in the ESC and the loci almost exhibit uniform spatial distribution in the chromosome (Figure 5A, blue lines in *Middle* and *Right* panels). During differentiation, the profile of radial density moves towards the interior of the chromosome along with specific re-positioning of active and inactive loci towards the surface and interior of the chromosome, respectively (Figure 5A and 5B). In further detail, for the regions with compartment A to B switching, the most probable state in the destined spatial distribution still allows a slight deviation from the interior (Figure 5C, *Left Most),* compared to the regions with stable compartment B to B (Figure 5C, *Right Most*), which undergoes the most significant spatial re-positioning towards the interior. For the regions with compartment B to A switching, the peak in the profile of radial density moves from the interior towards the surface of chromosome (Figure 5C, *Second Left),* but is still closer to the interior than the regions with stable compartment A to A (Figure 5C, *Second Right*). Therefore, we assume that the spatial re-positioning of chromosomal loci strongly depends on the compartment states during differentiation. The chromosomal regions maintaining the stable compartment states move more significantly in terms of the compartment A towards the surface of the chromosome and the compartment B towards the interior of the chromosome than the ones that have compartment A/B switching.

**Figure 5:**
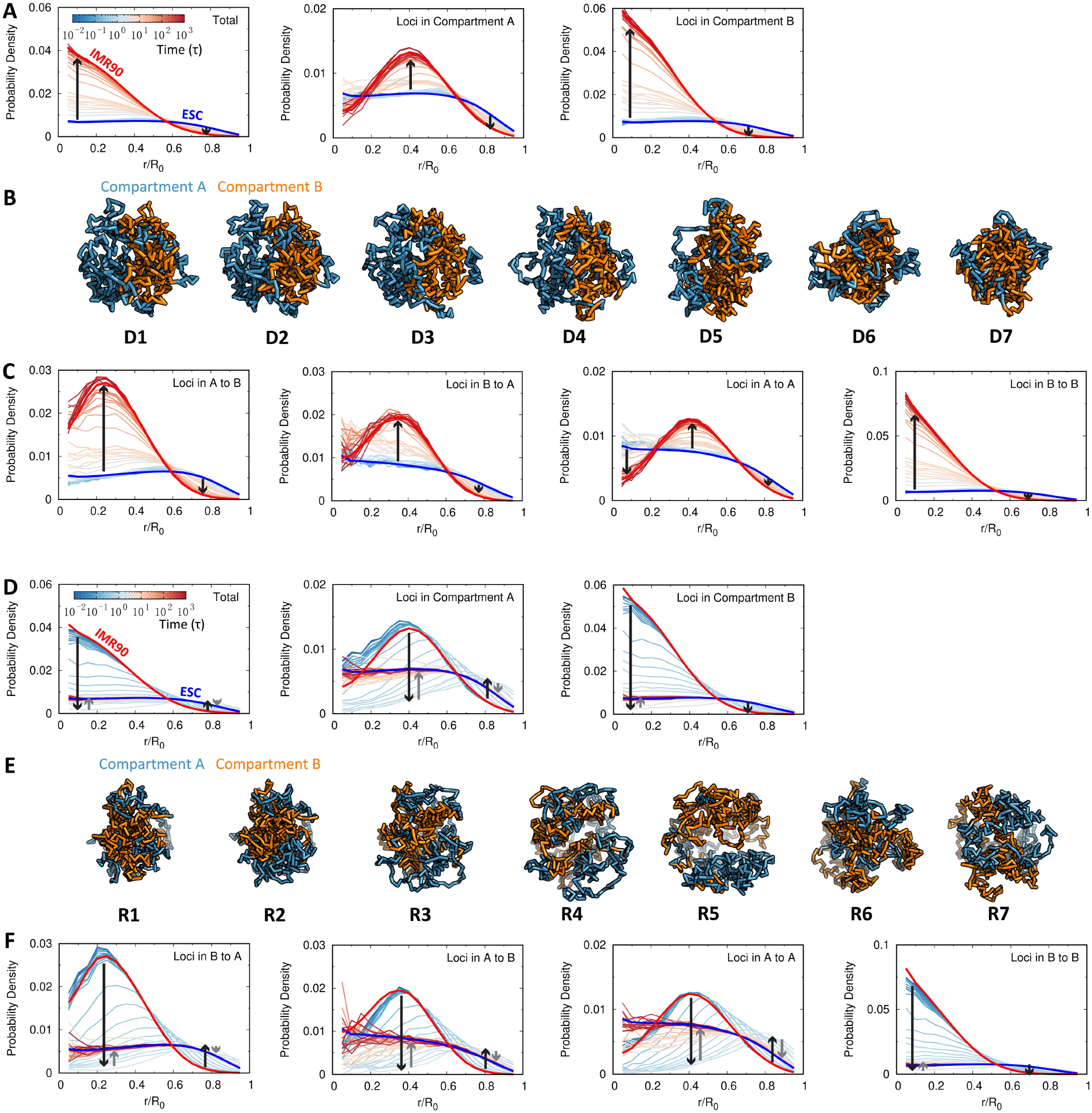
The change of the radial density in the chromosome during differentiation and reprogramming. (A) The change of the radial density profile for the whole loci *(Left),* loci in compartment A *(Middle)* and loci in compartment B *(Right)* in the chromosome during differentiation. The profile at each state is colored by the processing time in the logarithmic scale, while the ones at the ESC and IMR90 are colored in blue and red, respectively. (B) Structural illustrations of the chromosome with loci colored by compartment states during differentiation. (C) The change of the radial density profile for particular groups of loci in the chromosome during differentiation. The chromosomal loci are classified into 4 groups based on their associated compartment-switching patterns adapted during the differentiation, including dynamical switching of the compartment “A to B” and “B to A”, as well as the stable maintenance of the compartment “A to A” and “B to B” *(Left* to *Right).* The probability density is normalized by the number of loci involved. The lines with arrows illustrate the directions of changes in the radial density profile. (D-F) as (A-C) except for reprogramming. There are two types of lines with arrows in each figure for reprogramming. The black lines represent the expansion of the chromosome at the first stage, while the grey lines represent the compaction of the chromosome at the second stage.

On the other hand, we observed a completely different way of chromosome rearranging its spatial distribution during reprogramming, which is not the reversal of that in differentiation (Figure 5D). Notably, all loci in the chromosome segment, except the ones with stable compartment B to B, have to spatially move to the surface of the chromosome at the early stage of reprogramming, as an indication of chromosome overexpansion, followed by a slight re-positioning towards the interior of the chromosome (Figure 5E and 5F). Therefore, the whole process appears to be highly homogenous for chromosomal loci, independent of the compartment states of the chromosome during reprogramming. This eventually leads to nearly uniform spatial distribution of chromosomal loci in the ESC.

### 2.5 Bifurcation of chromosomal structural rearrangement paths during differentiation

We analyzed the experimental Hi-C data of four human early ESC-derived cell lineages, including trophoblast-like cell (TB), mesendoderm (ME), neural progenitor cell (NP) and mesenchymal stem cell (MS) [22]. As shown in Figure 6A, these cells represent the early ESC developmental stages and have deterministic pathways in differentiation to become either the extraembryo or to the three germ layers of the embryo [51]. We calculated the compartment profiles of the four cell lines from the experimental Hi-C data [22] and then incorporated them into the principal component analysis (PCA) on the trajectories of the chromosome compartment profiles evolving during differentiation in our simulations (Figure 6B). Interestingly, we found that except the ME, the other three experimental cell lines are located far from the ESC to the IMR90 differentiation pathway. This in general indicates that differentiation of the ESC to the IMR90 does not go through the TB, NP and MS cell lines at the chromosomal level. It is worth noting that the ME cell line is very close to the ESC [52] and there are also high similarities in Hi-C data between the ME and ESC, making the ME undistinguishable with the ESC in the PCA plot. Further clustering of the chromosome compartment profiles of the experimental cell lines along with the reduced 7 stages obtained from the differentiation simulations, shows that the ME, TB, NP cells are located at the early stage of differentiation, while the MS is close to the late differentiation process (Figure 6B, hierarchical clustering tree plot). This is consistent with the fact that the MS exhibits many characteristics of the cells at the relatively late developmental stages [51].

**Figure 6:**
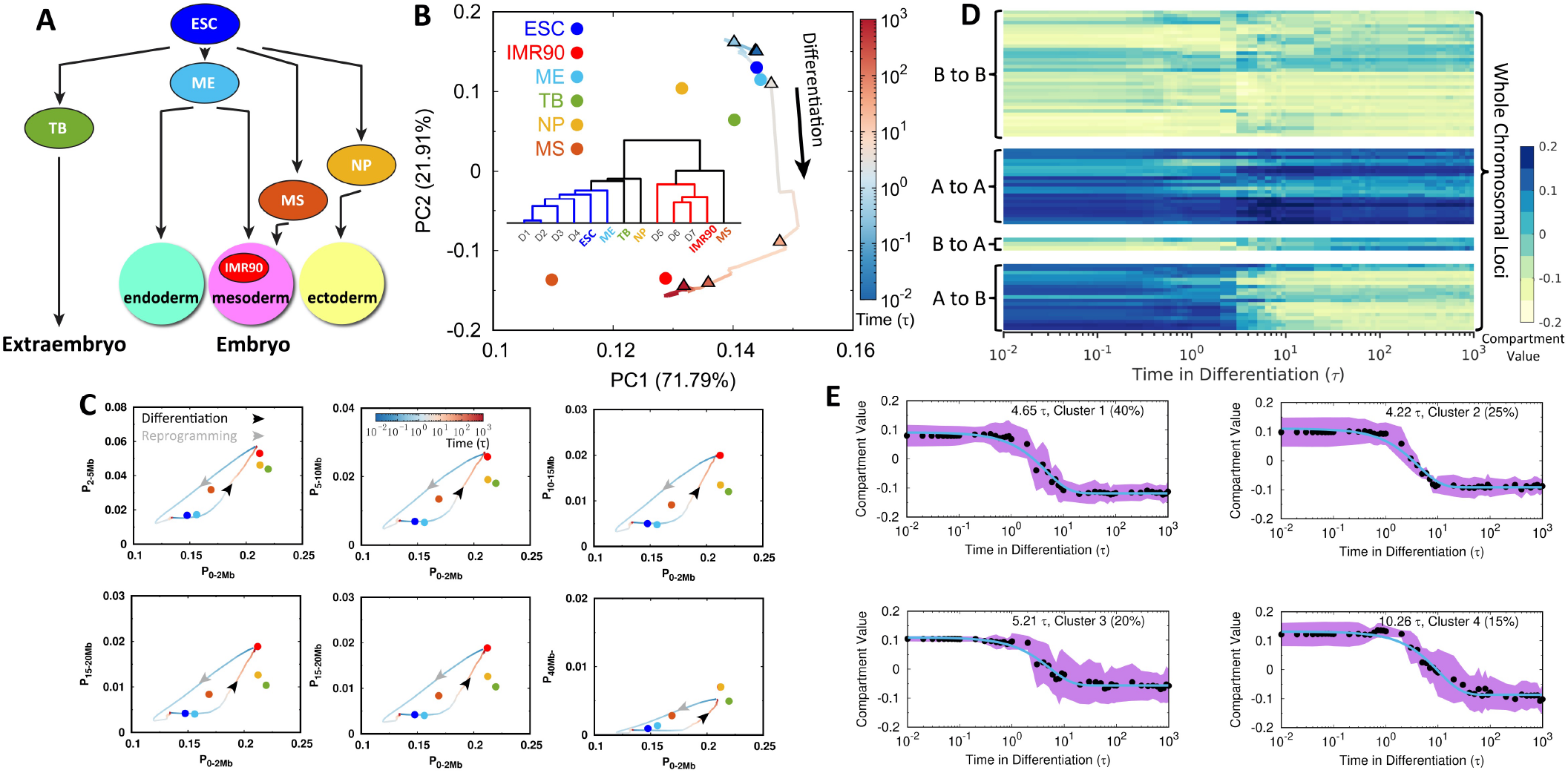
Bifurcation of the chromosomal structural rearrangement during differentiation. (A) Scheme illustrating the ESC differentiation to the extraembryo and three germ layers in the embryo along with the experimental cell lines that are used for comparisons in our study. (B) Principal component analysis (PCA) of chromosome compartment profiles during differentiation projected on the first two principal components (PCs) with populations indicated. Insert shows the hierarchical clustering tree of chromosome compartment profiles for the 7 stages in differentiation from the simulations and the experimental cell lines. (C) Change of the contact probability in the chromosome during differentiation and reprogramming varied by different contact ranges. Time is in logarithmic scale and arrows indicated the directions of differentiation (black) and reprogramming (grey). Experimental cell lines are calculated from Hi-C data indicated as points, in the same color scheme with those in (B). (D) Compartmental value of individual chromosomal locus changing with respect to differentiation time. Chromosomal loci are classified into 4 groups based on their types of compartment-switching in differentiation: A to B, B to A, A to A and B to B *(Bottom* to Top). (E) k-means clustering (k=4) of compartmental values changing with respect to time for loci that undergo compartment-switching from A to B, corresponding to the group at the bottom in (D). The averaged compartmental value of loci changing with time in each cluster is fitted to exponential function and the corresponding relaxation time is shown. The percentage is the population of the corresponding cluster. The standard deviation is shown with the shadow region.

Further projecting the pathways onto contact probability at various pairwise contact ranges of the chromosome clearly shows that the TB, NP and MS are neither on the pathways of the ESC differentiating to the IMR90, nor on the pathways of the IMR90 reprogramming to the ESC (Figure 6C). A similar observation can also be obtained by performing PCA on compartment profiles (Figure S11). Such non-overlapped chromosomal structural rearrangement in differentiation and reprogramming was observed in recent Hi-C experiments on mouse [53].

Since the compartment profiles of the chromosome change significantly during differentiation, we grouped the chromosomal loci based on the patterns that they adopt for compartment-switching and then examined the change of their compartmental values along the differentiation times (Figure 6D). There are 4 different groups of loci present with compartment-switching of A to B, B to A and stable compartment of A to A, B to B. Interestingly, for loci that do not undergo compartment-switching, they are not constantly stable, as apparent fluctuations were observed. This generally indicates that differentiation has extensive impacts spreading over the chromosome rather than only gradually switching the relevant compartments. As the compartment-switching of A to B contributes to the most significant chromosomal structural rearrangement that potentially impacts the genomic function and guides the direction of differentiation, we further investigated the kinetics of the compartment-switching of A to B (Figure 6E). We found that a large number of relevant loci undergo a similar timescale 4 ~ 5τ to perform the switching and a small number of loci is associated with a slower switching timescale ~ 10τ. Similar phenomena were found in the experiments where the compartment switches universally and gradually to the differentiated cells, while loci may be heterogeneous for participating in chromosomal structural rearrangement during differentiation [53].

## 3 Discussion

The landscape-switching model developed here and then applied to the cell differentiation and reprogramming processes produced the large-scale structural rearrangements occurring in a long human chromosome segment. We found that most of the TAD boundaries conserve during these two processes, in concordance with many experimental findings that TADs are conserved between different cell types [10, 22, 54]. Nevertheless, slight structural changes of TADs were observed with a rapid increase and decrease of TAD size during differentiation and reprogramming, as evidence to show that the chromosome structure and dynamics regulate the transcription activity during these two processes [55]. On the other hand, the relatively slow and notable changes in chromosome compartments were observed in terms of switching compartment A/B during differentiation and reprogramming, recapitulating the findings existing in Hi-C data [22] and the analysis that epigenomes differ significantly for various cell types [56, 51]. Interestingly, we found distinct scenarios for the chromosome to perform the compartment-switching during differentiation and reprogramming. Compartment-switching undergoes monotonic change during differentiation, though a small degree of fluctuations may be encountered due to cell stochasticity [57]. In contrast, during reprogramming, compartment profiles show aberrant changes in extensive regions, where compartmental values may be significantly different from the differentiated cell or the destined ESC.

By projecting the trajectories onto several order parameters, we obtained the chromosomal structural rearrangement pathways during differentiation and reprogramming. We attributed the aberrant changes in the compartment profiles during reprogramming to the over-expansion that occurs at both local and non-local contact ranges in the chromosome, though under different timescales. Such chromosome structure with spatial repositioning of the chromosomal loci universally to the border of chromosome territory (Figure 5D-F), is followed by slight compaction. Interestingly, the recent time series Hi-C experiment with a focus on the cell-cycle mitotic exit process identified a short-lived intermediate state at telophase, where the Hi-C properties cannot be derived from the combinations of those at the prometaphase and interphase [58]. This implies that the structure of the chromosome in such intermediate state does not lay in between the condensed cylinder in prometaphase or decondensed fractal globule in interphase [40]. Further analysis of the chromatin-protein binding experiments showed that the chromosome is free of SMC complexes at such intermediate state. In our recent study on the cell cycle, we established a correspondence of the over-expanded chromosome observed in the simulations to the intermediate state in the experiments [28]. The over-expanded chromosome topologically increases the accessible surface area for the loading of the SMC complexes, thus it may promote the chromosomal structural rearrangement in favor of an efficient mitotic exit process [59]. Likewise, we suggest a similar biological role of the overexpanded chromosome in facilitating the reprogramming process by opening the structure for protein binding. Different from cell differentiation, cell reprogramming is quite inefficient [60, 61]. From computational efforts, here we can propose a strategy to accelerate the reprogramming by opening the chromosome structure through chromosome remodeling, which has been found to play an important role in reprogramming of the somatic cells to pluripotency [62, 63].

The chromosomal structural rearrangement pathways observed in our simulations show high stochasticity and irreversibility for the differentiation and reprogramming processes. This finding is consistent with increasing experimental evidence that reprogramming proceeds in a way that deviates largely from the simple reversal of differentiation [64, 53]. Recent experiments interestingly showed that the mouse ESC can differentiate to the same destined state via multiple paths [65], as evidence of stochasticity. From a physical perspective, these facts underscore the prevalence of non-equilibrium and stochastic characteristics in cell differentiation and reprogramming, where the cell is constantly under intrinsic and extrinsic fluctuations from the underlying biochemical reactions involving a large number of molecules and the external stimuli in making cell-fate decision [66]. Therefore, the chromosomal structural rearrangements occur in a highly stochastic manner during differentiation and reprogramming. Still, the averaged irreversible pathways can be determined to some extent, driven by the destined states.

By developing a theoretical framework, we previously investigated the landscape and path of cell-fate decision-making processes based on the gene regulatory network [67, 68]. The resulting probability landscape provides a quantitative basis for the original pictorial Waddington’s epigenetic landscape [21]. This can be used to guide the dynamics of cell development. Furthermore, it was shown that the cell-fate decisionmaking processes are additionally determined by the curl force (flux), which originates from the broken detailed balance in the non-equilibrium dynamics [69]. The detailed balance breaking leads to the non-equilibrium thermodynamic cost for maintaining the functional cell states, supplied by the underlying molecular processes (e. g., ATP hydrolysis). As the epigenetic information is encoded in genome architecture [70, 50], we can suggest a landscape based on our simulation findings at the chromosomal level (Figure 7). We showed that from the chromosome structural perspective, the differentiation paths from the ESC to the IMR90 are well-defined and bifurcate at a very early stage from the paths differentiating to the other cells, consistent with our theoretical predictions [67]. The pathways do not necessarily follow the steepest descent gradient paths (along the valleys) because of the non-equilibrium effects. Thus, they may bypass the MS, which is a metastable attractor on the landscape. The deviation is controlled by the directions and magnitudes of the flux, which can drive the developmental paths in practice to go through distinct differentiated intermediate states in reaching the destination [65]. As the efficiency of reprogramming from the somatic cells to the iPSCs is generally lower than 1% [60, 61], the terminally differentiated cells are stable and likely located at the minima of the landscape and reprogramming thus externally requires the introductions of reprogramming factors into the system. In addition, reprogramming undergoes different paths that do not overlap with the differentiation ones, due to the non-equilibrium effects, in line with the theoretical predictions [67]. Chromosomes on the differentiation and reprogramming paths located at the same level as the same layer on the landscape should be structurally different. This is also observed experimentally [53], when the mouse B cell was reprogrammed to a state which resembles the epiblast, but is very different at the genomic and genetic level in terms of Hi-C and RNA-seq data.

**Figure 7:**
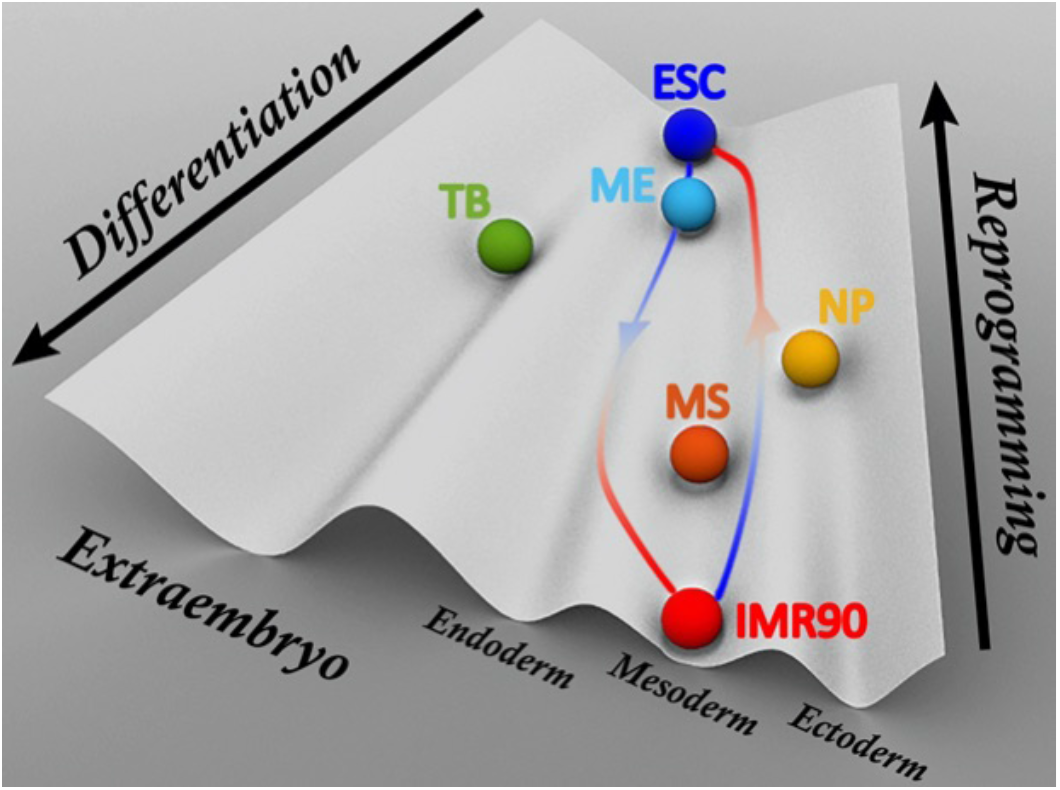
A schematic Waddington landscape for cell differentiation and reprogramming gained from a perspective of chromosomal structural rearrangement. The balls are attractors on the landscape and colored in the same scheme as Figure 6 to represent the cell lines used in our study. The paths are shown for illustrating the differentiation (*Left*) and reprogramming (*Right*) processes.

As evidenced by the experiments [71, 72], the gene regulatory network that determines the cell development process shows multi-stability, corresponding to different cell fates. Besides, the transitions between these cell fates in the cell development have been found to occur abruptly, similar to a switch between different gene expression states [73, 74, 75]. Recent theoretical studies showed that the simple circuitry of bi-stable switches between two gene expression states, which correspond to distinct cell fates, is able to capture many characteristics of the cell development in reality [76, 77, 78, 79]. Overall, our landscape-switching model is in accordance with the switching between the bi-stable cell fates in cell development. However, the model has two prominent limitations worth noting. First, cell development has been simplified as a bi-stable system, while the intermediate cell states, often recognized as the multipotent progenitor cells in differentiation, may exist during the process. Second, the model implements the instantaneous switch to trigger the transition between different cell fates, whereas the real biological responses in both cell differentiation and reprogramming should occur with finite reaction rates. The most plausible solution for overcoming these limitations is to add more intermediate cell fates into the model when the experimental data are available, so the model will eventually have the stepwise switches between the multi-stable cell states during cell differentiation and reprogramming. Besides, our model does not consider the cell cycle dynamics, due to the lack of experimental Hi-C data and the knowledge of the quantitative relation between the cell cycle and the cell development. In reality, there are usually many cell cycles during cell differentiation and reprogramming. Since we are mainly interested in the development process in this study, for simplicity we focused on the slow developmental process while treating the fast cell-cycle dynamics as an averaged background. Therefore, the model provided a prediction of the chromosome structural evolution during cell differentiation and reprogramming at the interphase, which accounts for most of the time in the cell cycle. Further improvements in implementing the cellcycle dynamics into cell development should be considered in the future.

Our computational predictions can be potentially assessed via the targeted experiments. In this regard, it would be highly useful to apply the time-series Hi-C experiments, which were recently developed and applied to the cell cycle process [80, 81, 58, 82], to obtain the time-evolving Hi-C maps during differentiation and reprogramming. Cell differentiation and reprogramming depend on both the intrinsic and extrinsic factors, so the processes may vary under different conditions. The switching implementation imposes an instantaneous change of the conditions to the desired ones, so our observations may correspond to the extremes, where the transitions between cell fates are strongly favored. With rapid developments in improving the reprogramming efficiency [83, 84, 85, 86], we anticipate that the predictions present here can be tested by the experiments in the future.

## 4 Conclusions

In summary, we explicitly considered the non-equilibrium dynamics and investigated chromosomal structural rearrangement during cell differentiation and reprogramming through a landscape approach. We simplified the non-equilibrium effects as an instantaneous energy excitation followed by the subsequent relaxation, so the kinetics was driven by the landscape-switching, which resonates with the cell-fate switching in multi-stable cell development. Our approach provides a physical and structural evolution picture of the cell-fate decision-making processes for differentiation and reprogramming through the chromosome structural dynamics. The predictions made in this study are in need of assessments by future experiments. Our model can be extended to gain insights into the chromosome pathological disorganization in cancer cells and cell senescence.

## 5 Model and Simulation Section

### 5.1 Hi-C data processing

We used human embryonic stem cell (ESC) and human fetal lung fibroblast cell (IMR90), a terminally differentiated cell, as the two start/end points to investigate the differentiation and reprogramming processes. Hi-C data of these two cell lines were measured by Dixon et al. [10] and are publicly available on GEO with accession number GSE35156. Hi-C data for the four cell lines (TB, ME, NP and MS) during the early ESC development used for comparisons in our study can be downloaded on GEO with accession number GSE52457 [22]. Hi-C data were handled by the Hi-C Pro software through the standard pipelines [87]. To best minimize the effects of the experimental errors existing in Hi-C experiments, all the replicas for each cell line were collected and used for generating the Hi-C contact map at a resolution of 100kb.

### 5.2 Maximum entropy principle simulations

We used a generic polymer model as the basic background [36], which is made up of bonded potentials to maintain the chain connection and soft-core nonbonded potentials to allow chain crossing as an outcome of topoisomerase [43]. Topoisomerases, which are found abundant in cell nucleus [88], show wide participation in most of the reactions that involve doublestranded DNA, including transcription and recombination [89], which are essential for cell differentiation and reprogramming. As noted elsewhere [33], the underlying kinetic behaviors of the chromosome loci are not affected by the soft-core potentials when the chromosome is within one cell state. Therefore, we applied the soft-core potentials throughout the simulations to keep the effects of the topoisomerase being constantly active.

In order to best reproduce the Hi-C map for the ESC and IMR90, we implemented the experimental restraints into the generic polymer model by means of maximum entropy principle [31, 32]. The potential of the polymer model for the ESC or IMR90 can be written as:

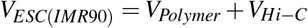

where *V_Polymer_* is the potential of the generic polymer model and the restraint potential from the Hi-C data (*V_Hi-C_*) is linearly added with *V_Polymer_* as requested by the maximum entropy principle [90]. Hi-C maps were normalized to be contact probability maps *f_i,j_*, where *i* and *j* are chromosomal loci, by assuming the adjacent beads always form contacts *f*_*i,i*±1_ ≡ 1 [31]. Therefore *V_Hi-C_* has an expression:

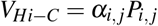

where *P_i,j_* is the calculated contact probability using step function and *α_i, j_* is the parameter that is determined by the maximum entropy principle through multiple rounds of iteration to ultimately match *P_i,j_* to *f_i,j_*. Details can be found in previous studies [31, 32].

### 5.3 Landscape-switching model

A prominent outcome of the maximum entropy principle simulation is the potential *V_ESC(IMR90)_*, which can generate a chromosome ensemble in matching the Hi-C map and reproduce the kinetic essence of chromosome dynamics in experiments [34, 33]. We observed the sub-diffusivity of the genomic loci in the chromosome at both the ESC and IMR90 (Figure S6). The diffusion exponents, which are characterized in the relation of mean square displace *(MSD)* and time *t* with *MSD ~ t^β^*, are found to be *β* ~ 0.4, in line with previous experimental observations [91, 92].

In our previous work on the cell cycle [28], we developed an energy excitation-relaxation landscape-switching model to approximately and explicitly include the non-equilibrium effects in simulations to describe the chromosome conformational transition between the interphase and mitotic phase for the cell-cycle dynamics. Here, we extended our model to the ESC differentiation and the IMR90 reprogramming. In practice, the non-equilibrium simulations were performed in three steps:

i. Chromosome was initially simulated under the potential *V_ESC_* (preparing for differentiation) or *V_IMR90_* (preparing for reprogramming);
ii. A sudden switch of the potential from *V_ESC_* to *V_IMR90_* (triggered to differentiation) or *V_IMR90_* to *V_ESC_* (triggered to reprogramming) was implemented;
iii. Chromosome was relaxed under the new potential *V_IMR90_* (differentiation) or *V_ESC_* (reprogramming) for a long period of time.

### 5.4 Identifications of compartments and boundaries of TADs

To identify the compartments, we first calculated the enhanced contact probability map (Figure S9), which is the observed versus expected contact map, at a reduced resolution level (1Mb) [9,93]. Then we used Iterative Correction and Eigenvector decomposition (ICE) method to normalize the enhanced contact probability map [94] and subsequently convert it into a Pearson correlation matrix. Finally, we followed the previous suggestion to perform the principal component analysis (PCA) on this matrix, and the first PC (PC1) was assigned as the compartment profiles [9]. The direction of the PC1 values is arbitrary, and we set the positive and negative PC1 values with gene density (positive to gene-rich and negative to gene-poor). This is done on the IMR90 data. Then, the direction of the PC1 values during differentiation and reprogramming was determined according to the correlation coefficient with the PC1 values of the IMR90 Hi-C data.

The boundaries of TADs were identified by the insulation score proposed by Crane et al. [44]. We used the same size of sliding square (500×500kb) within the original application [44] to calculate the insulation score. The local minima of the insulation score represent the areas of high local insulation, which may correspond to the boundaries of TADs. We then used a delta vector that measures the slope of the insulation vector to quantitatively determine the boundaries of TADs [44]. Practically, the delta vector with crossing the horizontal 0 and difference between the local left maximum and local right minimum higher than 0.1 signified the boundary of a TAD. To calculate the boundary matching population, we allowed 100kb noise error in accounting as an exact matching. We found that the average numbers of beads in forming the TADs for the chromosome focused in our study identified from the Hi-C maps are about 9 and 10 in the ESC and IMR90, respectively. Forming the TADs by such relatively low numbers of beads could be contributed by the thermal fluctuations, which can transiently induce the contacts formed by the beads close in sequence. Therefore, we evaluated the TAD formation in our model for the chromosome ensemble at the ESC and IMR90. This was undertaken by comparing to the polymer ensemble generated by the simulations with applying only the generic polymer potential *V_Polymer_*. We found that the insulation score of the polymer ensemble constantly remains zero, corresponding to no TAD formation (Figure S5). The result indicated that the TADs are robustly formed in our chromosome model under the Hi-C constraint potentials *V_ESCiIMR90;_*.

### 5.5 Statistical Analysis

The hierarchical clustering on the chromosome ensemble generated by the maximum entropy principle simulations was performed on the randomly selected 2400 chromosome structures. The cut-off criteria were chosen to be the up-limit of structural difference for chromosome conformational diffusion (Figure S3 and S4). By these criteria, chromosomes within one cluster are structurally similar, as shown by the contact maps. Two structures from each cluster, whose population is higher than 0.3%, were chosen. These structures were used as the initial structures for the landscape-switching simulations. There are 354 and 226 simulation trajectories for differentiation and reprogramming, respectively. All the trajectories were collected and used for calculating the contact probability, compartment profiles, insulation scores, radial density and the averaged pathways. The changes of TAD size and compartmental values for A to B switching with the time were presented as mean ± standard deviation.

## Supporting Information

Supporting Information is available on bioRxiv.org.

## Acknowledgements

We thank Shufan Ma for help in preparing the figures. We acknowledge the support from the National Science Foundation PHY-76066. The authors would like to thank Stony Brook Research Computing and Cyberinfrastructure, and the Institute for Advanced Computational Science at Stony Brook University for access to the high-performance SeaWulf computing system, which was made possible by a $1.4M National Science Foundation grant (#1531492).

## Table of Contents

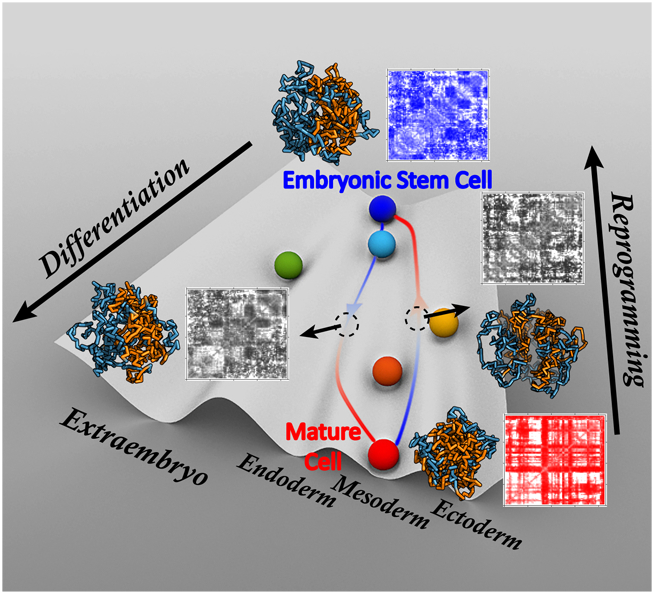

**ToC text:** Chromosomes are the structural scaffolds for gene functions, which determine the cell developmental process. A non-equilibrium landscape model is developed to identify the chromosomal structural and dynamical origins of cell differentiation and reprogramming. The irreversible pathways in differentiation and reprogramming show distinct dynamical rearrangement for the chromosome structure to facilitate the formation of specific gene expression patterns and associated functions.

**ToC keyword:** Irreversible chromosome dynamics

## Notes

### Competing Interest Statement

The authors have declared no competing interest.

