## Supplementary Material for "Microscopic Chromosomal Structural and Dynamical Origin of Cell Differentiation and Reprogramming"

for

### 1 Additional materials and methods

We used Gromacs (version 4.5.7) [1] with PLUMED (version 2.5.0) [2] to undertake all the simulations in our study. The polymer was coarse-grained into a "beads-on-a-string" model and initially governed by the generic polymer potential $V_{Polymer}$ [3]. Soft-core repulsive potentials between the non-bonded beads were used to mimic the effects of topoisomerase [4]. Reduced units were used with time and length units denoted as $\tau$ and $\sigma$, respectively. $\sigma$ can be regarded as the size of the bead, and it corresponds to the length of the pseudo-bond that is described by the finitely extensible nonlinear elastic (FENE) potential [5] in the generic polymer model potential $V_{Polymer}$. A spheric confinement was additionally applied to ensure the volume fraction of the chromosome to be 0.1 [6]. Further details of the potential $V_{Polymer}$ can be found here [7, 8] and our previous work [9].

The maximum entropy principle simulations were performed routinely for chromosome dynamics in the ESC and IMR90, respectively, following the standard protocols proposed in previous studies [7, 8]. The aim of the maximum entropy principle is to rationally implement the experimental Hi-C data to the polymer model in order to generate a chromosome ensemble, which is as close as possible to the prior knowledge implies [10]. As requested by the maximum entropy principle, the potential of chromosome in the ESC or IMR90 is linear in the observables (Hi-C data):

$$V_{ESC(IMR90)} = V_{Polymer} + V_{Hi-C}$$

$V_{Hi-C}$ is experimentally restraint potential in the form of a sum of a function of pairwise contact probability as a matter of maximum entropy principle [10]:

$$V_{Hi-C} = \sum_{i,j} \alpha_{i,j} P_{i,j}$$

, where $P_{i,j}$ is the contact probability and $\alpha_{i,j}$ is the parameter for genomic loci $i$ and $j$. The process was performed with multiple iterations to minimize the differences between simulation contact probability $P_{i,j}$ and Hi-C contact probability $f_{i,j}$ by adapting the parameters $\alpha_{i,j}$ regularly. After a reasonable convergence criterion was met [7], the chromosome ensemble was generated by collecting the frames in the trajectories. We found quantitative matching of Hi-C maps from simulations to experiments (Figure S1 and S2). We observed high correlations on many of the aspects regarding TADs and compartments between simulations and experiments (Figures S1 and S2). This evidence validates the chromosome ensemble generated by our simulations for the ESC and IMR90.

Next, we investigated the kinetics of the chromosome dynamics under the Hi-C restraint potential $V_{ESC(IMR90)}$. We calculated the mean square displacement (MSD) along with the time for the loci in the chromosome (Figure S6). We found the diffusion exponents $\beta$ in the relation of $MSD \sim t^{\beta}$ at both the ESC and IMR90 are about 0.4, which signifies the sub-diffusivity of the loci in chromosome dynamics and is also in line with experimental observations [11, 12]. With available experiments, we further mapped the simulation units to the physical units. We approximately regarded the 100kb bead in our model as the 30-nm fiber, which gives the Kuhn length unit $\sigma \approx 150nm$. The experiments on measuring the diffusion of the chromosome in the HeLa cell show that the MSD at $0.5s$ is in the range of 0.01-0.015$\mu m^2$ [12]. Our simulation data eventually have $\tau \approx$ 3.3-4.2$s$ (by mapping to the ESC) or $\tau \approx$ 2.5-3.6$s$ (by mapping to the IMR90). It is worth noting that due to the essence of the landscape-switching model, the timescale estimated within one cell state may not be directly linked to the real reaction rates of the cell differentiation and reprogramming processes.

The landscape-switching simulations started from the structures obtained from the abovementioned approaches. To investigate the chromosomal structural rearrangements during differentiation (reprogramming), first, the chromosome was simulated under $V_{ESC}$ ($V_{IMR90}$) for a duration of $5000\tau$. Then, a sudden switch of the landscape $V_{ESC} \rightarrow V_{IMR90}$ ($V_{IMR90} \rightarrow V_{ESC}$) was implemented. Finally, chromosome was simulated under $V_{IMR90}$ ($V_{ESC}$) for a duration of $10000\tau$. In practice, we found that $1000\tau$ is sufficient for a full relaxation of chromosome under the new potential (landscape), so we only present the data till $1000\tau$ in order to have a better visualization.

#### 2 Additional figures

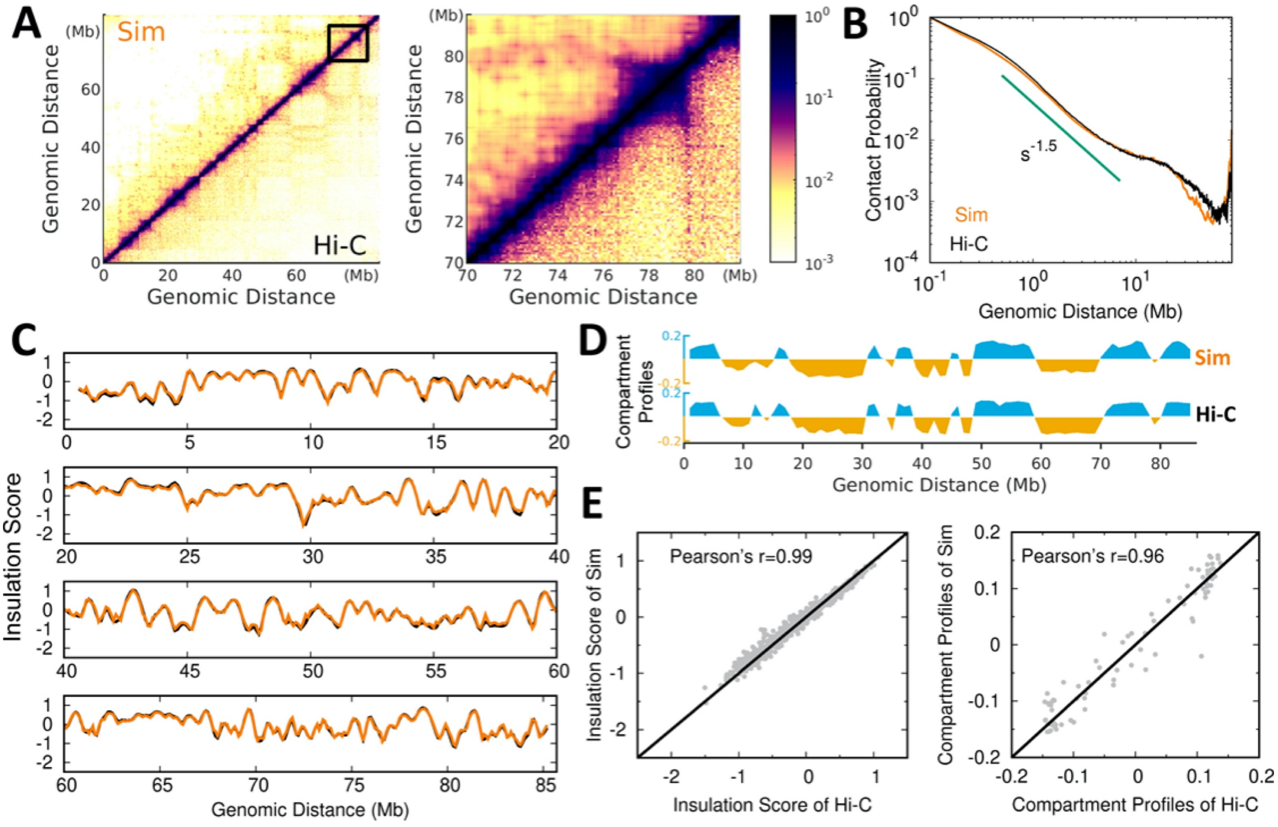


Figure S1: Comparisons of simulations and Hi-C data in the ESC. (A) Hi-C contact maps of chromosome from maximum entropy principle simulations and experiments at global (*Left*) and local (*Right*) scales. (B) Contact probability versus genomic distance in chromosome for simulations and Hi-C data with a slope of -1.5 observed. (C) Insulation score of chromosome obtained by simulations and Hi-C data. (D) Compartment profiles of chromosome obtained by simulations and Hi-C data. (E) Correlations of insulation score (*Left*) and compartment profiles (*Right*) between simulations and Hi-C data.

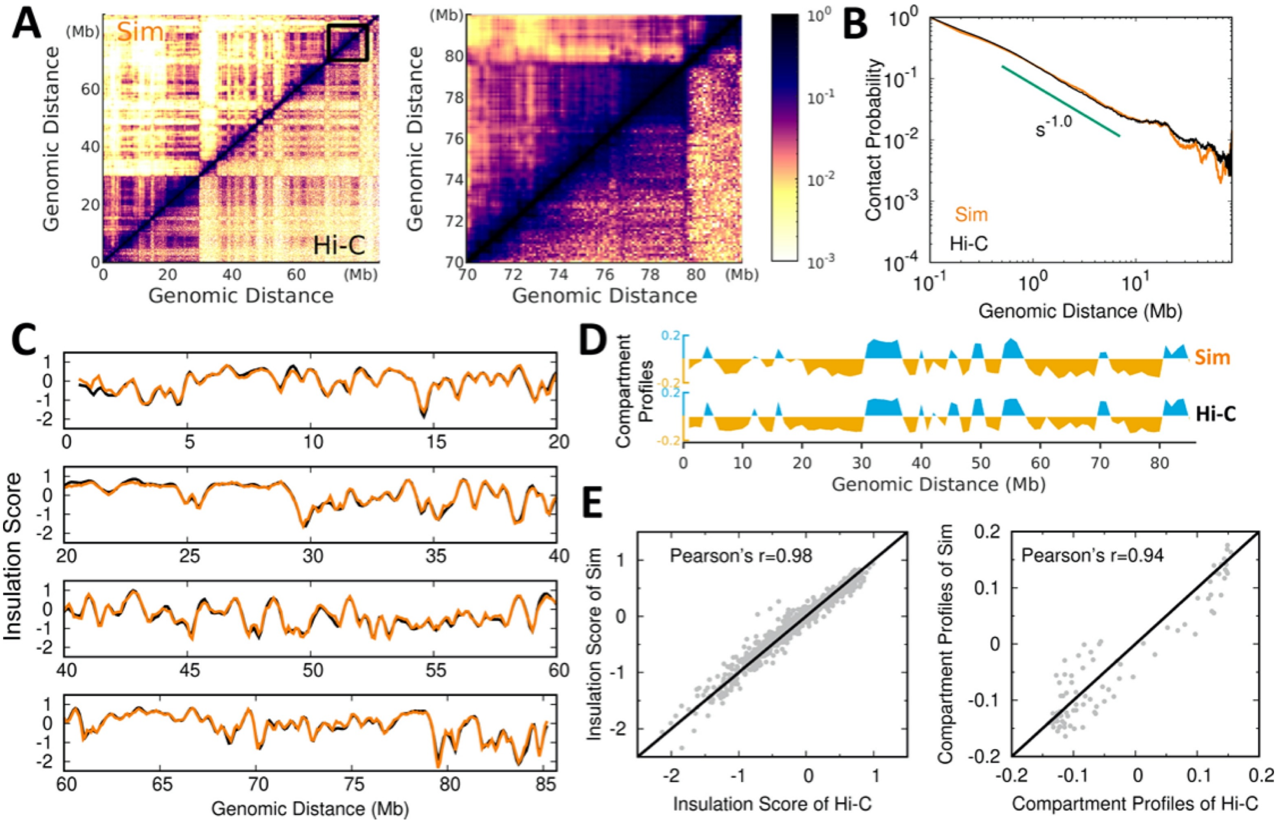


Figure S2: Comparisons of simulations and Hi-C data in the IMR90. (A) Hi-C contact maps of chromosome from maximum entropy principle simulations and experiments at global (*Left*) and local (*Right*) scales. (B) Contact probability versus genomic distance in chromosome for simulations and Hi-C data with a slope of -1.0 observed. (C) Insulation score of chromosome obtained by simulations and Hi-C data. (D) Compartment profiles of chromosome obtained by simulations and Hi-C data. (E) Correlations of insulation score (*Left*) and compartment profiles (*Right*) between simulations and Hi-C data.

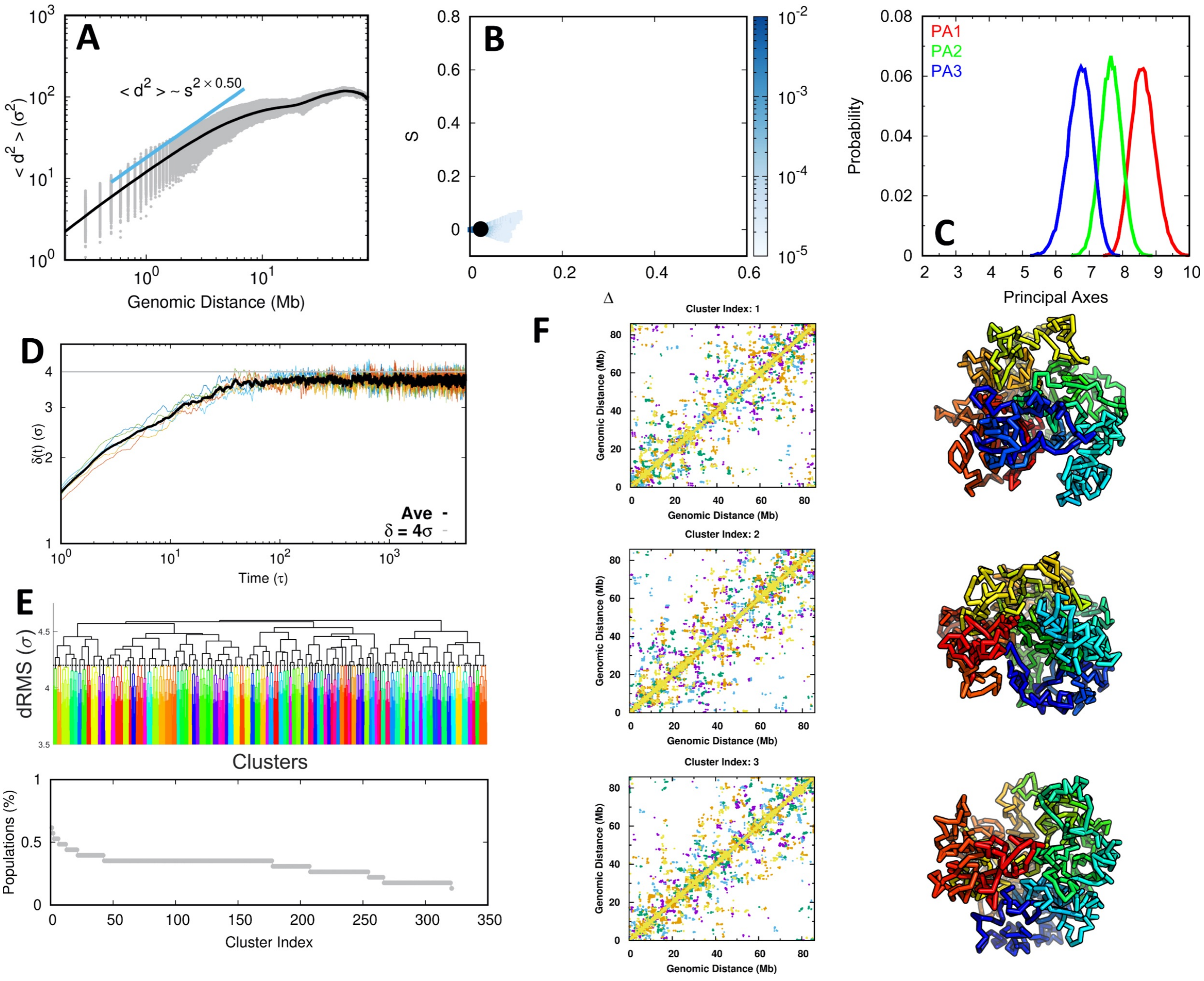


Figure S3: Chromosome ensemble in the ESC. (A) Contact distance ($d$) versus genomic distance in the chromosome. (B) The probability distributions of aspheric parameters of the chromosome. $\Delta$ and $s$ are calculated using the inertia tensor [13]. Deviation of $\Delta$ from 0 (the value corresponding to a sphere) gives an indication of the extent of anisotropy. Negative values of $S$ correspond to oblate shapes and positive values to prolate shapes. (C) The probability distribution of configurational extension on three principal axes of the chromosome. (D) The time evolution of average root mean square distance ($d_{rms}$) between every genomic pair in chromosome at the time $t$ relative to its initial value: $\delta(t) = \sum_{i,j} d_{rms}(i,j,t)/N_{pairs}$, where $N_{pairs}$ is the number of summed pairs. The maximum of $\delta(t)$ is close to $4\sigma$. (E) The hierarchical clustering of chromosome shown as a dendrogram (*Top*) and the populations of the cluster (*Bottom*). Cut-off distance $4\sigma$ was applied. (F) The top 3 most populated chromosome clusters. Each is shown with a mixed contact map (*Left*), which contains 5 structures within the cluster, and one representative structure (*Right*).

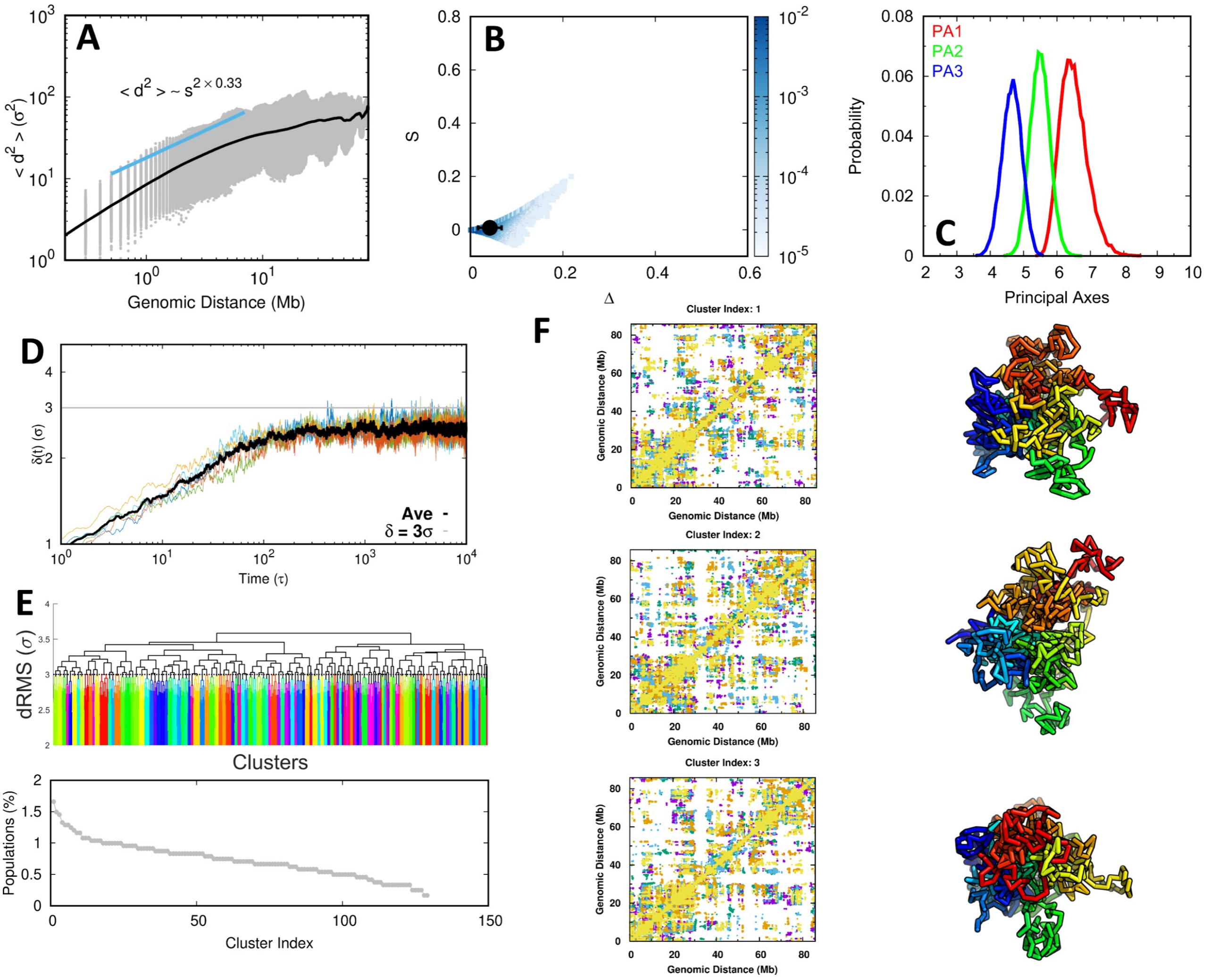


Figure S4: Chromosome ensemble in the IMR90. (A) Contact distance ($d$) versus genomic distance in the chromosome. (B) The probability distributions of aspheric parameters of the chromosome. (C) The probability distribution of configurational extension on three principal axes of the chromosome. (D) The time evolution of $\delta(t)$. The maximum of $\delta(t)$ is close to $3\sigma$. (E) The hierarchical clustering of chromosome shown as a dendrogram (*Top*) and the populations of the cluster (*Bottom*). Cut-off distance $3\sigma$ was applied. (F) The top 3 most populated chromosome clusters. Each is shown with a mixed contact map (*Left*), which contains 5 structures within the cluster, and one representative structure (*Right*).

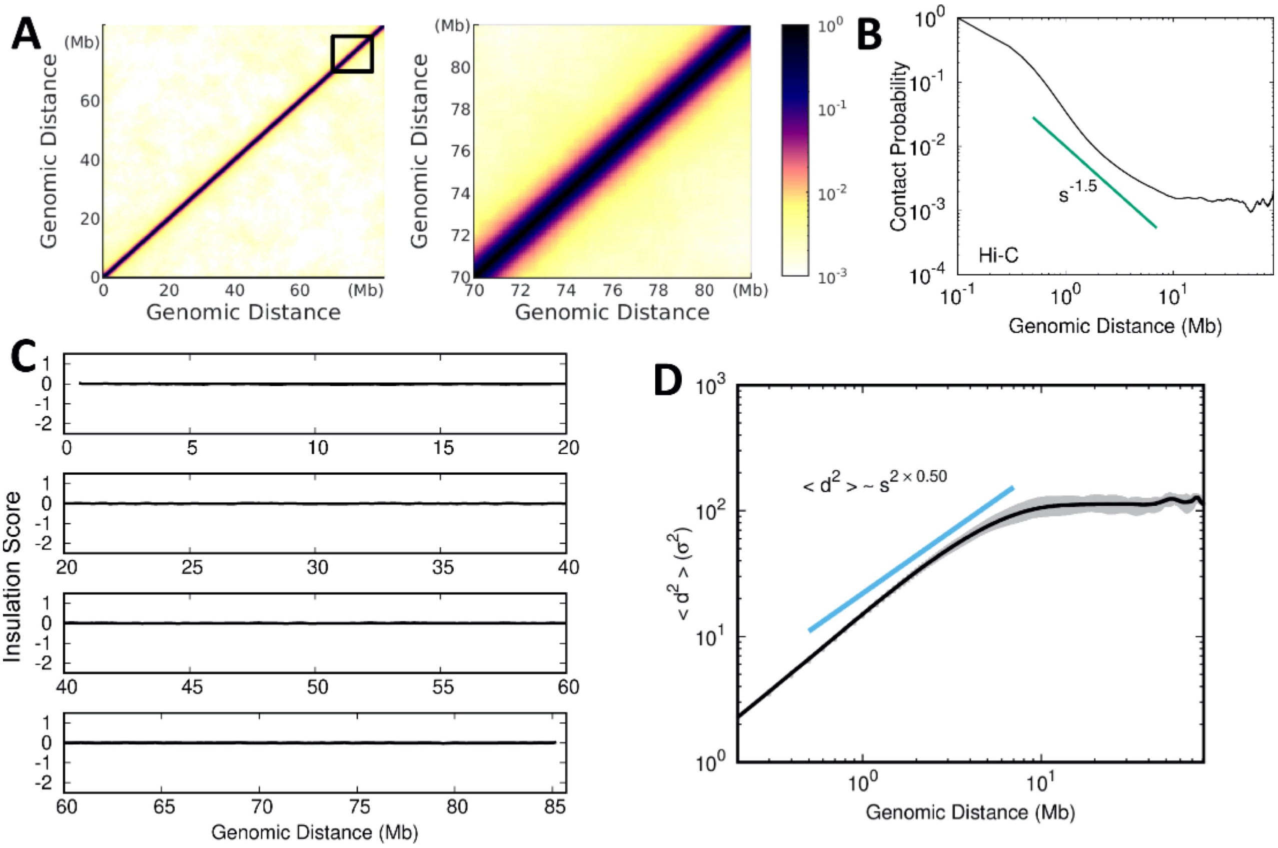


Figure S5: Chromosome ensemble generated by $V_{Polymer}$. (A) Hi-C contact maps of the polymer at global (*Left*) and local (*Right*) scales. (B) Contact probability versus genomic distance with a slope of -1.5 observed. (C) Insulation score. (D) Contact distance (d) versus genomic distance.

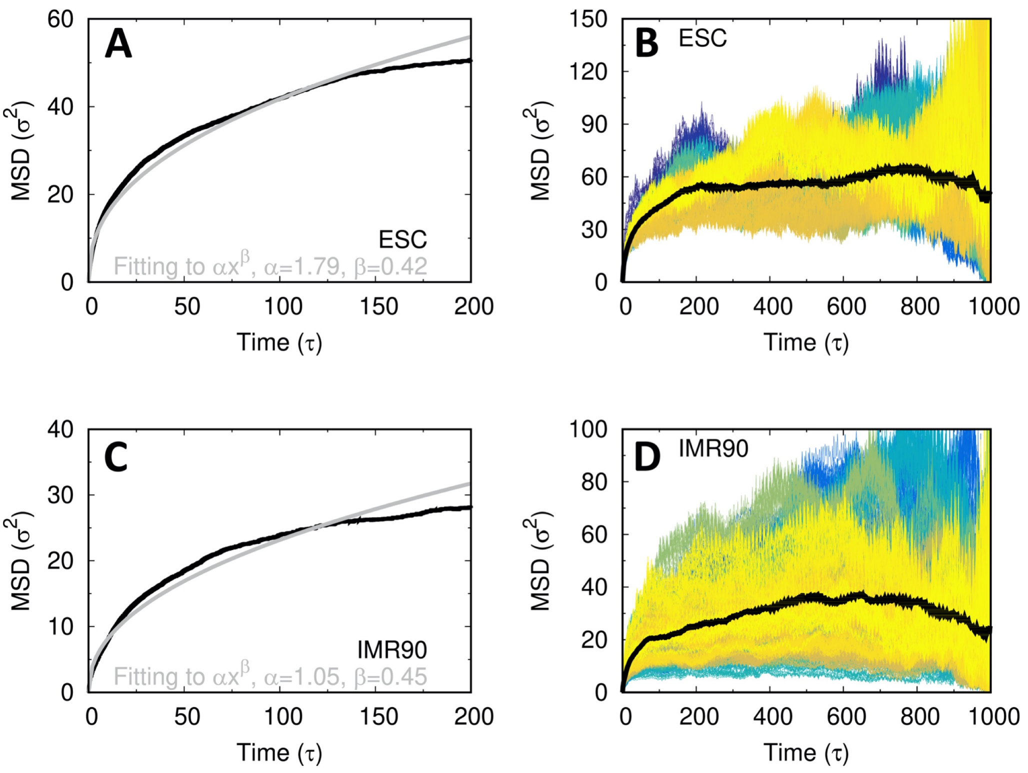


Figure S6: Chromosome diffusion dynamics in the ESC and IMR90. (A) Fitting of MSD to the power-law function in the ESC. MSD is averaged from 5 independent simulations. (B) MSD of all the individual genomic loci in the ESC obtained from one simulation with the average shown as the black line. (C) and (D) as (A) and (B) but for IMR90.

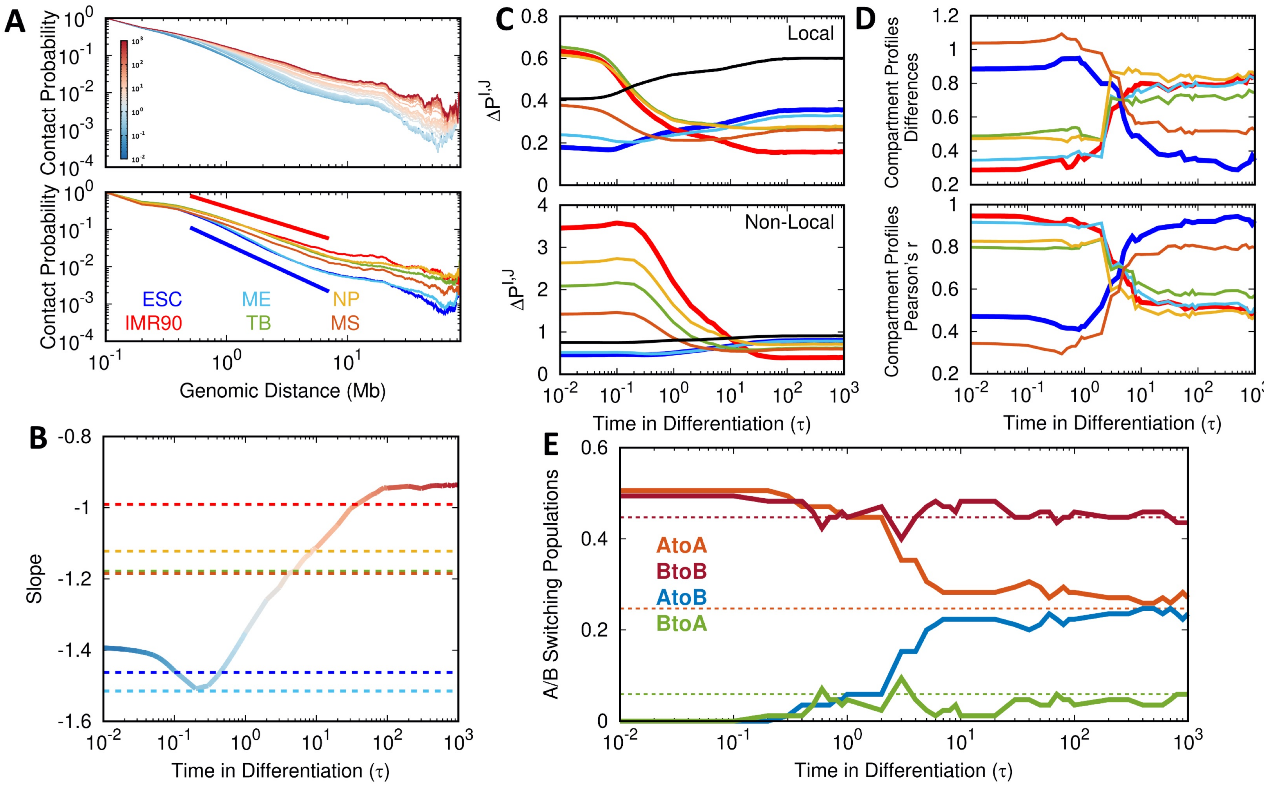


Figure S7: Chromosomal structural rearrangements during differentiation. (A) The evolution of contact probability $P(s)$ versus genomic distance $s$ in chromosome during differentiation (*Top*) and comparisons of $P(s)$ for the Hi-C data of the cell lines used in our study (*Bottom*). (B) The evolution of the slope ($a$) in the relation $P(s) \sim s^a$, the dashed lines dictate the values for the Hi-C data of the cell lines used in our study. (C) The evolution of contact probability differences in chromosome between the differentiation time and the cell lines used in our study at local (*Top*) and non-local (*Bottom*) contact ranges. (D) The evolution of compartment profiles differences (*Top*) and correlations (*Bottom*) in chromosome between the differentiation time and the cell lines used in our study. (E) The evolution of compartment switching populations in the chromosome. The dashed lines dictate the destined values.

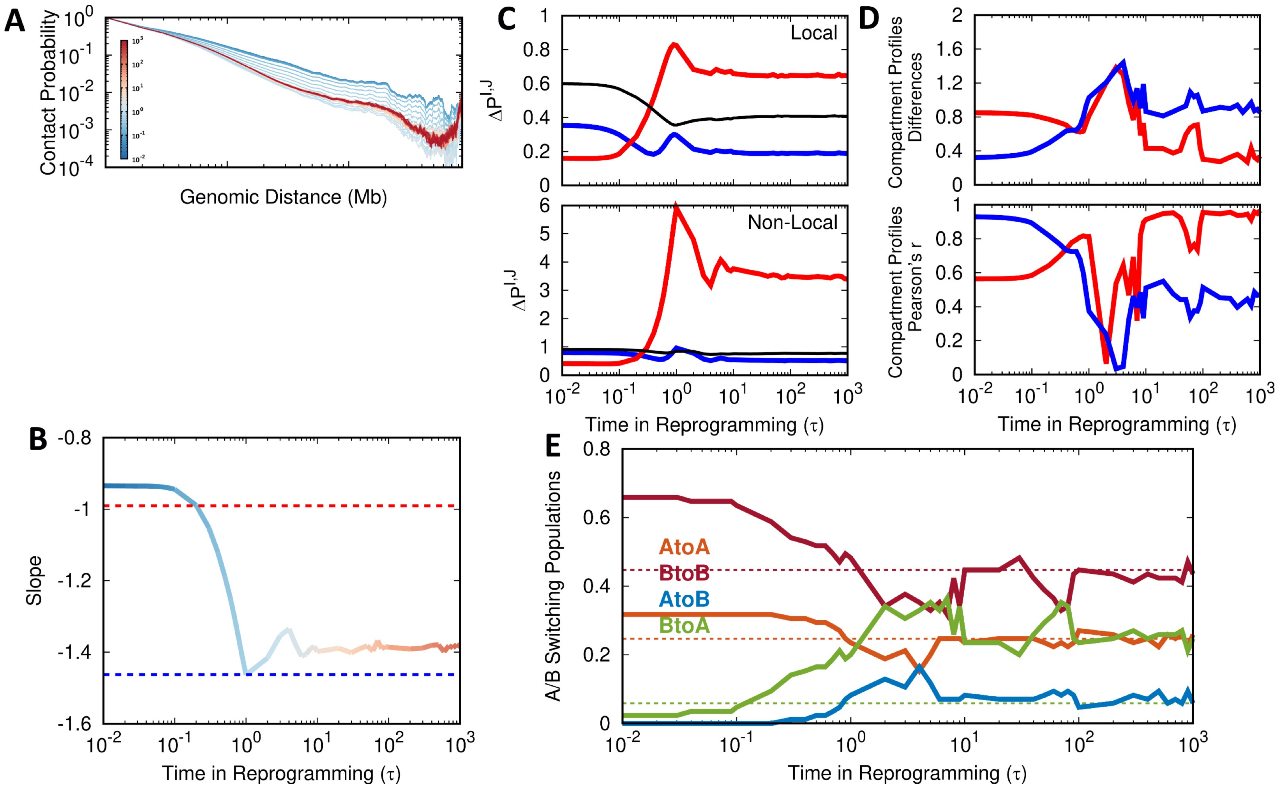


Figure S8: Chromosomal structural rearrangements during reprogramming. (A) The evolution of contact probability $P(s)$ versus genomic distance $s$ in chromosome during reprogramming. (B) The evolution of the slope $(a)$ in the relation $P(s) \sim s^a$, the dashed lines dictate the values for ESC and IMR90. (C) The evolution of contact probability differences in chromosome between the reprogramming time and the ESC and the IMR90 at local (*Top*) and non-local (*Bottom*) contact ranges. (D) The evolution of compartment profiles differences (*Top*) and correlations (*Bottom*) in chromosome between the differentiation time and ESC and IMR90. (E) The evolution of compartment switching populations in the chromosome. The dashed lines dictate the destined values.

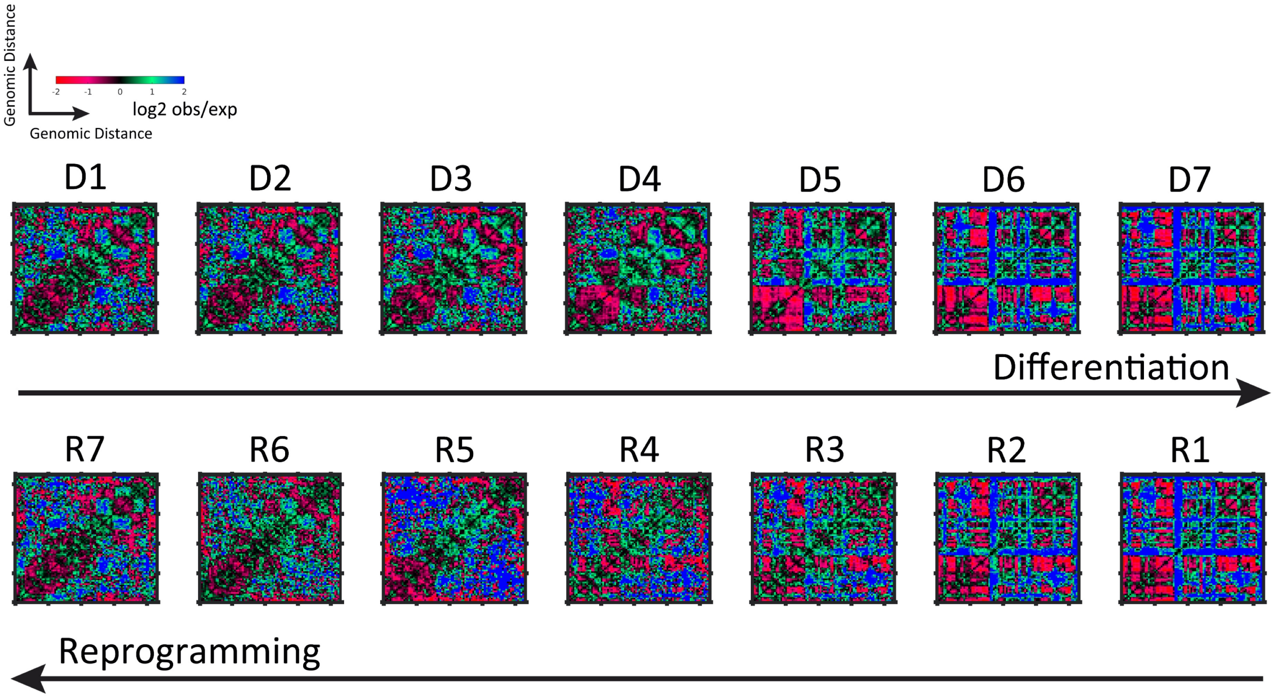


Figure S9: Enhanced contact probability matrices of chromosome for the 7 stages in differentiation and reprogramming. The enhanced contact probability is calculated by observed versus expected contact probability at a reduced resolution of 1Mb. The expected matrix is calculated as the mean contact probability at a given genomic distance, as suggested by Nagano et al. [14].

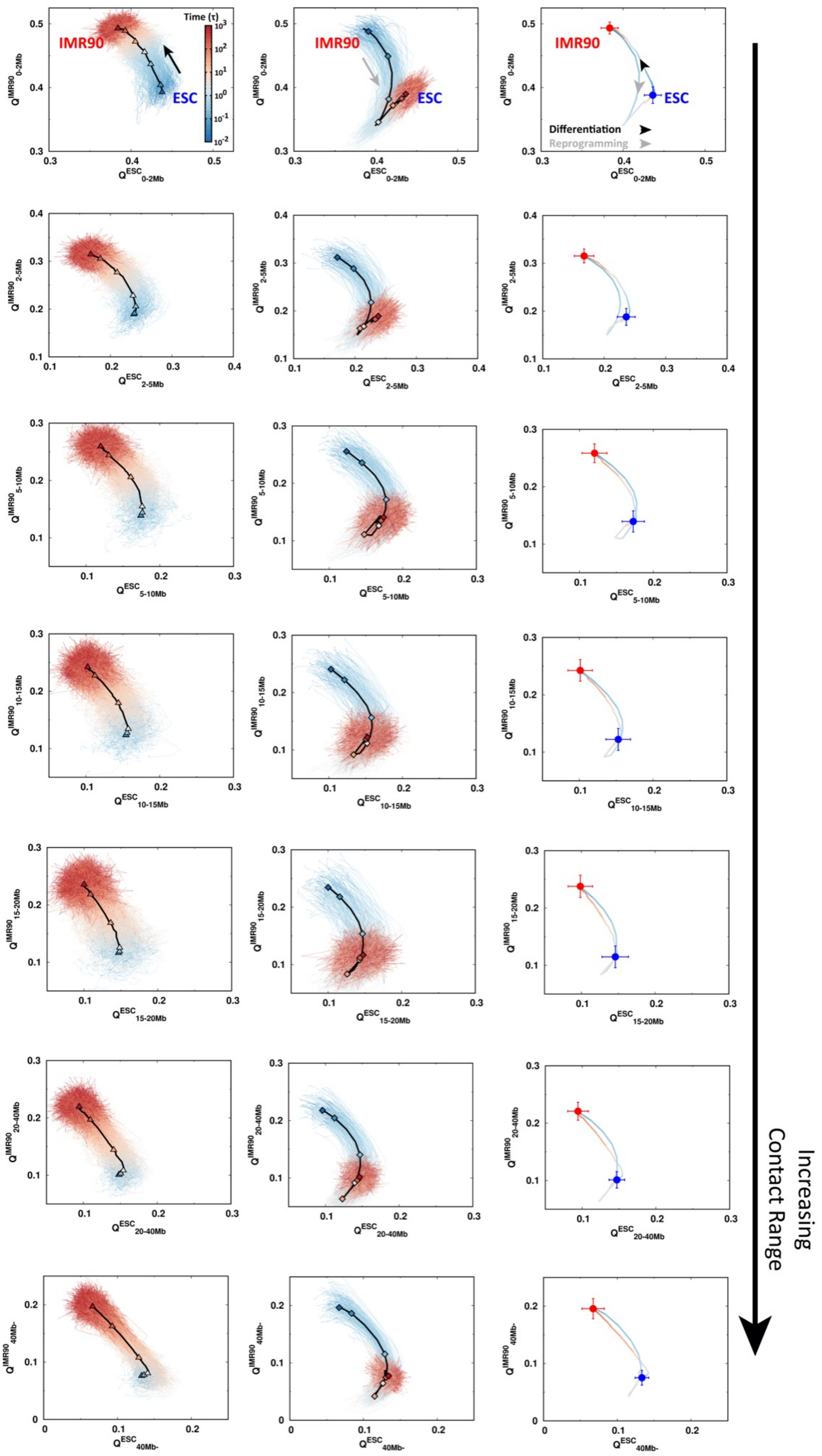


Figure S10: Pathways of chromosomal structural rearrangements projected onto $Q$ varied by different contact ranges during differentiation (*Left*) and reprogramming (*Middle*), as well as their averages (*Right*).

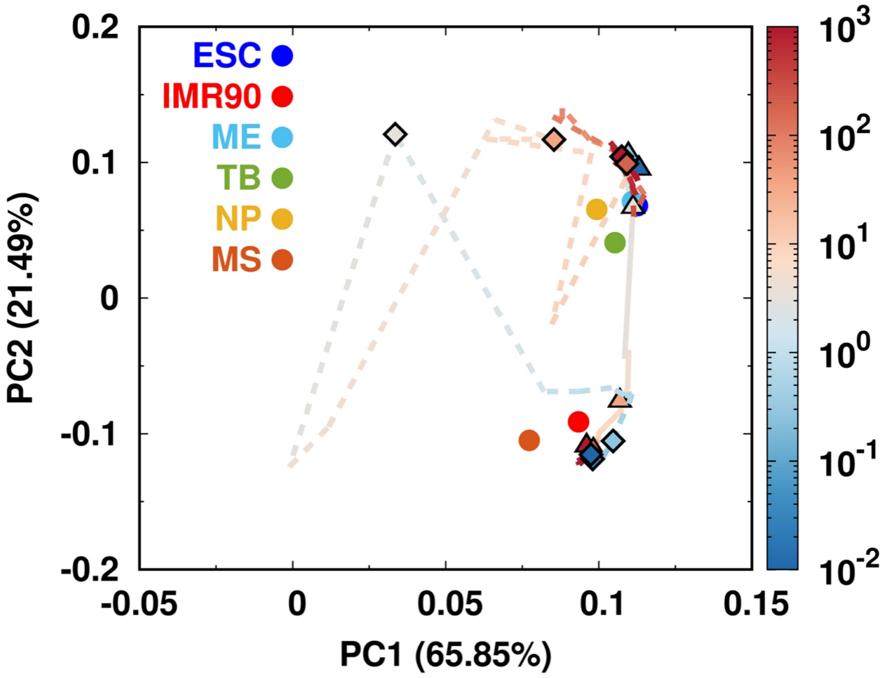


Figure S11: The PCA of compartment profiles of chromosome during differentiation (*Solid line*) and reprogramming (*Dashed line*). The differentiation stages "D1-D7" are shown in triangles and reprogramming stages "R1-R7" are shown in diamonds. The cell lines used in our study are indicated.
